# Evaluation of parameters affecting performance and reliability of machine learning-based antibiotic susceptibility testing from whole genome sequencing data

**DOI:** 10.1101/607127

**Authors:** Allison L. Hicks, Nicole Wheeler, Leonor Sánchez-Busó, Jennifer L. Rakeman, Simon R. Harris, Yonatan H. Grad

## Abstract

Prediction of antibiotic resistance phenotypes from whole genome sequencing data by machine learning methods has been proposed as a promising platform for the development of sequence-based diagnostics. However, there has been no systematic evaluation of factors that may influence performance of such models, how they might apply to and vary across clinical populations, and what the implications might be in the clinical setting. Here, we performed a meta-analysis of seven large *Neisseria gonorrhoeae* datasets, as well as *Klebsiella pneumoniae* and *Acinetobacter baumannii* datasets, with whole genome sequence data and antibiotic susceptibility phenotypes using set covering machine classification, random forest classification, and random forest regression models to predict resistance phenotypes from genotype. We demonstrate how model performance varies by drug, dataset, resistance metric, and species, reflecting the complexities of generating clinically relevant conclusions from machine learning-derived models. Our findings underscore the importance of incorporating relevant biological and epidemiological knowledge into model design and assessment and suggest that doing so can inform tailored modeling for individual drugs, pathogens, and clinical populations. We further suggest that continued comprehensive sampling and incorporation of up-to-date whole genome sequence data, resistance phenotypes, and treatment outcome data into model training will be crucial to the clinical utility and sustainability of machine learning-based molecular diagnostics.

**Author Summary:** Machine learning-based prediction of antibiotic resistance from bacterial genome sequences represents a promising tool to rapidly determine the antibiotic susceptibility profile of clinical isolates and reduce the morbidity and mortality resulting from inappropriate and ineffective treatment. However, while there has been much focus on demonstrating the diagnostic potential of these modeling approaches, there has been little assessment of potential caveats and prerequisites associated with implementing predictive models of drug resistance in the clinical setting. Our results highlight significant biological and technical challenges facing the application of machine learning-based prediction of antibiotic resistance as a diagnostic tool. By outlining specific factors affecting model performance, our findings provide a framework for future work on modeling drug resistance and underscore the necessity of continued comprehensive sampling and reporting of treatment outcome data for building reliable and sustainable diagnostics.

## Introduction

At least 700,000 deaths annually can be attributed to antimicrobial resistant (AMR) infections, and, without intervention, the annual AMR-associated mortality is estimated to climb to 10 million in the next 35 years (1). As most patients are still treated based on empirical diagnosis rather than confirmation of the causal agent or its drug susceptibility profile, development of improved, rapid diagnostics enabling tailored therapy represents a clear actionable intervention (1). The Cepheid GeneXpert MTB/RIF assay, for example, has been widely adopted for rapid point-of-care detection of *Mycobacterium tuberculosis* (TB) and rifampicin (RIF) resistance (2), and the SpeeDx ResistancePlus GC assay used to detect both *Neisseria gonorrhoeae* and ciprofloxacin (CIP) susceptibility was recently approved for marketing as an *in vitro* diagnostic in Europe.

Molecular assays offer improved speed compared to gold-standard phenotypic tests and are of particular interest because of their promise of high accuracy for the prediction of AMR phenotype based on genotype (2, 3). Approaches for predicting resistance phenotypes from genetic features include direct association (*i.e.*, using the presence or absence of genetic variants known to be associated with resistance to infer a resistance phenotype) and the application of predictive models derived from machine learning (ML) algorithms. Direct association approaches can offer simple, inexpensive, and often highly accurate resistance assays for some drugs/species (2) and may even provide more reliable predictions of resistance phenotype than phenotypic testing (4–6). However, these approaches are limited by the availability of well-curated and up-to-date panels of resistance variants, as well as the diversity and complexity of resistance mechanisms. ML strategies can facilitate modeling of more complex, diverse, and/or under-characterized resistance mechanisms, thus outperforming direct association for many drugs/species (7–9). With the increasing speed and decreasing cost of sequencing and computation, ML approaches can be applied to genome-wide feature sets (8, 10–18), ideally obviating the need for comprehensive *a priori* knowledge of resistance loci.

While prediction of antibiotic resistance phenotypes from ML-derived models based on genomic features has become increasingly prominent as a promising diagnostic tool (8, 11–15, 17), there has been no systematic evaluation of factors that may influence performance of such models and their implications in the clinical setting. The extent to which ML model accuracy varies by antibiotic is unclear, as is the impact of sampling bias on model performance. It is further unclear what the most relevant resistance metric (*i.e.*, minimum inhibitory concentration [MIC] or categorical report of susceptibility) for such a diagnostic might be and how amenable different species might be to genotype-to-phenotype modeling of antibiotic resistance.

We used set covering machine (SCM) (19) and random forest (RF) (20) classification as well as RF regression algorithms to build and test predictive models with seven gonococcal datasets for which whole genome sequences (WGS) and ciprofloxacin (CIP) and azithromycin (AZM) MICs were available. AZM is currently part of the recommended treatment regimen for gonococcal infections, and with the development of resistance diagnostics, CIP may represent a viable treatment option (21–23). While the majority of CIP resistance in gonococci can be attributed to *gyrA* mutations, AZM resistance is associated with more diverse and complex resistance mechanisms (23, 24), offering an opportunity to evaluate ML methods across drugs with distinct pathways to resistance. The range of datasets and sampling frames enables assessment of sampling bias on model reliability. Further, the availability of MICs, as well as distinct European Committee on Antibiotic Susceptibility Testing (EUCAST) and Clinical and Laboratory Standards Institute (CLSI) breakpoints, for these drugs allows for evaluation of predictive models based on different resistance metrics. Finally, extension of these analyses to *Klebsiella pneumoniae* and *Acinetobacter baumannii* datasets for which WGS and CIP MICs were available allows for assessment of model performance for the same drug in species with open pangenomes (25, 26), which may be more difficult to model given the increased genomic diversity and potential resistance mechanism diversity and complexity (47).

Our results demonstrate that using ML to predict antibiotic resistance phenotypes from WGS data yields variable results across drugs, datasets, resistance metrics, and species. While more comprehensive assessment of different methods will be required to build the most accurate and reliable models, we suggest that tailored modeling for individual drugs, species, and clinical populations may be necessary to successfully leverage these ML-based approaches as diagnostic tools. We further suggest that continuing surveillance, isolate collection, and reporting of WGS, MIC phenotypes, and treatment outcomes will be crucial to the sustainability of any such molecular diagnostics.

## Results

### Accuracy of ML-based prediction of resistance phenotypes varies by antibiotic

Given the distinct MIC distributions and distinct pathways to resistance for CIP and AZM in gonococci, these two drugs enable evaluation of drug-specific performance of ML-based resistance prediction models. CIP MICs in surveys of clinical gonococcal isolates are bimodally distributed, with the majority of isolates having MICs well above or below the non-susceptibility (NS) breakpoints, while the majority of reported AZM MICs in gonococci are closer to the NS breakpoints (https://mic.eucast.org/Eucast2). These trends were recapitulated in the gonococcal isolates assessed here (**Fig 1a-b**). Further, the vast majority of CIP resistance in gonococci observed to date is explained by mutations in *gyrA* and *parC* and has spread predominantly through clonal expansion, generally resulting in MICs ≥ 1 μg/mL (23, 27). In contrast, AZM resistance in gonococci has arisen many times *de novo* through multiple pathways, many of which remain under-characterized and are associated with lower-level resistance (23, 27, 28). As expected, the GyrA S91F mutation alone predicts NS to CIP by both EUCAST and CLSI breakpoints in the aggregate gonococcal dataset assessed here with ≥98% sensitivity and ≥99% specificity (**Table S1**). AZM NS showed lower values for these metrics, indicating it was not as well explained by known resistance variants, with extensive contributions from uncharacterized mechanisms and/or multifactorial interactions (**Table S2**).

**Figure 1.**
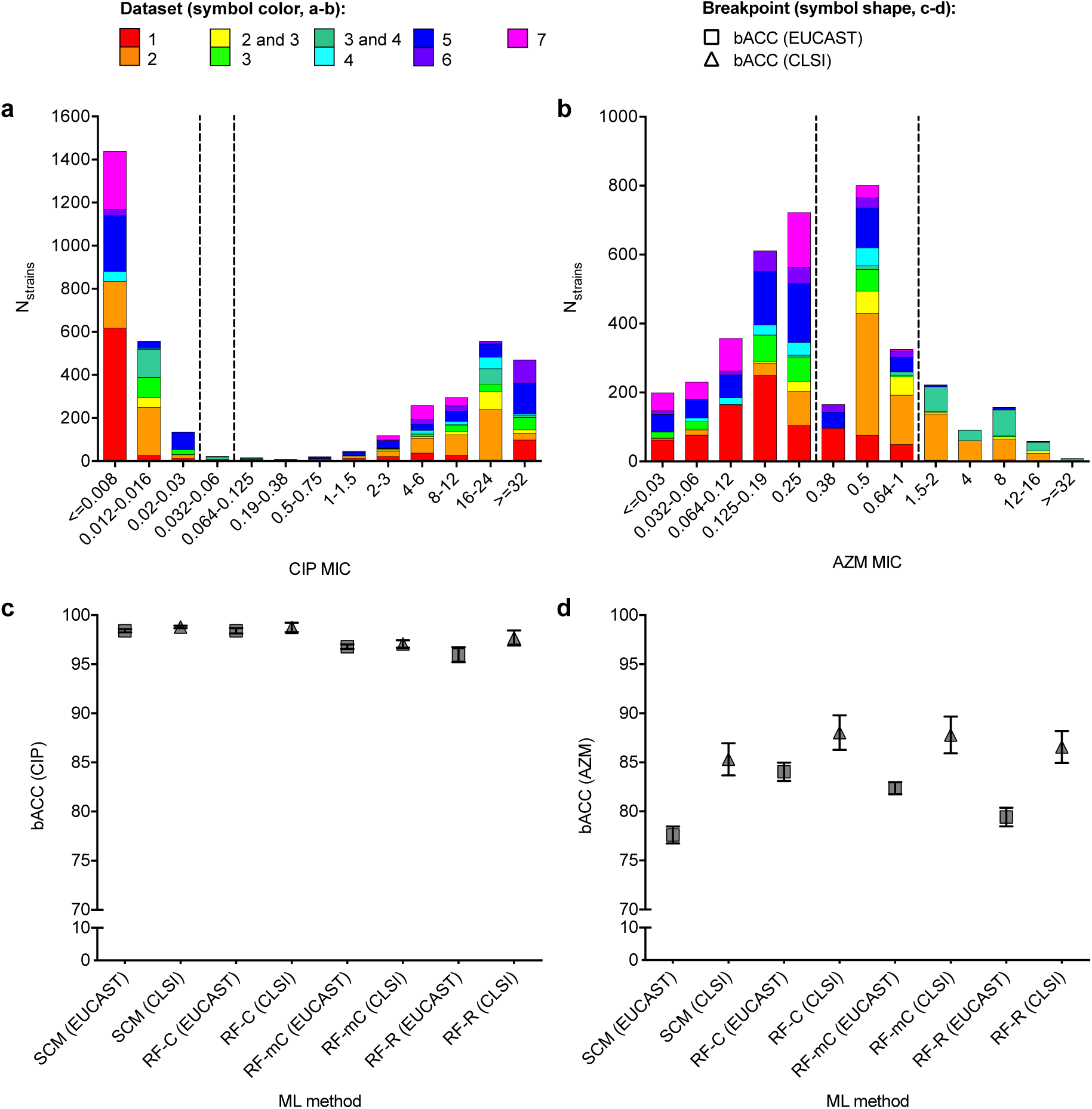
Differential performance of machine learning-based prediction models for ciprofloxacin and azithromycin resistance in gonococci. Histograms showing the distributions of **(a)** ciprofloxacin (CIP) and **(b)** azithromycin (AZM) minimum inhibitory concentrations (MICs) in the gonococcal isolates assessed here. Bar color indicates the study or studies associated with the isolates. Dashed lines indicate the **(a)** EUCAST and CLSI breakpoints for non-susceptibility (NS, >0.03 μg/mL and >0.06 μg/mL, respectively) for CIP and the **(b)** EUCAST and CLSI breakpoints for non-susceptibility (>0.25 μg/mL and >1 μg/mL, respectively) for AZM. Note that there was some overlap in strains from the US between datasets 2 and 3 and in strains from Canada between datasets 3 and 4; such strains are indicated in **(a)** and **(b)** as belonging to datasets 2 and 3 and 3 and 4, respectively. Mean balanced accuracy (bACC) with 95% confidence intervals of predictive models for **(c)** CIP NS and **(d)** AZM NS trained and tested on the aggregate gonococcal dataset. Symbol colors in (**a-b**) indicate the datasets from which the training and testing sets were derived. Symbol shapes in (**c-d**) indicate the NS breakpoint. SCM, set covering machine; RF-C, random forest classification; RF-mC, random forest multi-class classification; RF-R, random forest regression.

We next trained and evaluated ML-based predictive models for CIP and AZM resistance in gonococci (**Table S3**). By all ML methods and breakpoints, CIP NS was predicted with significantly higher balanced accuracy (bACC) than AZM NS in the aggregate gonococcal dataset (*P <* 0.0001, **Fig 1c-d**, **Tables S4-S5**): CIP NS was predicted with mean bACC ≥93% across all methods, breakpoints, and datasets, whereas mean bACC for AZM NS classification ranged from 57% to 94% (**Tables S4-S5**). Variation in model performance across antibiotics has been attributed to different proportions of susceptible (S) and NS isolates (7, 14, 15); however, by the EUCAST breakpoints, the aggregate gonococcal dataset as well as some of the individual datasets had nearly identical proportions of CIP and AZM susceptible and non-susceptible isolates, demonstrating that variable representation of S and NS isolates alone cannot explain reduced performance of AZM models compared to CIP.

We tested whether the poorer performance for AZM may be attributable to the large fraction of isolates with MICs around the breakpoint. Removing strains with AZM MICs that were ≤2 doubling dilutions of the NS breakpoints from the aggregate gonococcal dataset (**Table S6**) yielded AZM MIC distributions similar to those of CIP (**Fig S1a-b**). Analysis of this restricted dataset resulted in higher performance of SCM and RF AZM NS classifiers compared to those trained and tested on the full aggregate gonococcal dataset (**Fig S1c**). However, bACC of AZM classifiers trained and tested on the restricted datasets was still significantly lower than bACC of the CIP NS classifiers (*P* < 0.0001 and *P* < 0.003 for classifiers based on the EUCAST and CLSI breakpoints, respectively), suggesting that both MIC distribution and additional drug-specific factors can influence performance of resistance classifiers.

### Sampling bias in training and testing data skews resistance model performance

The diversity of resistance mechanisms for AZM in gonococci offers an opportunity to evaluate the effects of sampling bias on model performance. The sampling frames for the seven gonococcal datasets ranged geographically from citywide to international and temporally from a single year to >20 years, and several datasets were enriched for AZM resistance (11, 29) (**Table 1**). The distributions of both AZM MICs and known resistance mechanisms across datasets (**Fig 1b, Table S2**) and the variable performance of AZM resistance models across datasets (**Table S5**) suggest that AZM resistance mechanisms are differentially distributed across the sampled clinical populations. Further, the higher performance of many SCM and RF-based AZM classifiers on training data compared to test sets (**Table S5**) suggests that potentially due to a lack of signal, AZM models are incorporating substantial noise or confounding factors, which may be population-specific. To assess the impact of sampling on model reliability, the performance of RF classifiers in prediction of AZM NS phenotypes were compared across multiple training and testing sets. These include classifiers trained on subsamples of isolates from a single dataset, classifiers trained on the aggregate gonococcal dataset, and classifiers trained on the aggregate gonococcal dataset excluding isolates from the same dataset as the testing set (**Table S6**). Given the low representation of AZM NS strains by the CLSI breakpoint in many datasets, these analyses were only performed using the EUCAST breakpoint.

**Table 1.** Summary of datasets.

| Species | Dataset | SRA Study ID/Reference | N<sub>samples</sub> | Temporal range | Geographic range | Sampling approach |
| --- | --- | --- | --- | --- | --- | --- |
| *N. gonorrhoeae* | 1 | ERP011192 | 886 | 2011-2015 | New York, NY (US) | Survey from citywide clinics |
|  | 2 | ERP008891, ERP001405, ERP000144 (23) | 1102 | 2000-2013 | National (US) | Survey from nationwide clinics; male patients only; |
|  |  |  |  |  |  | enriched for CFX resistance |
|  | 3 | SRP065041, ERP000144, ERP001405, ERP008891, SRP072971 (29) | 671 | 2004-2014 | International (UK, Canada, US) | Surveys from Brighton, UK (60) and nationwide sites in Canada (30, 61) and the US (23, 62); Canadian samples enriched for CRO and AZM resistance; US samples enriched for CFX resistance; US samples from male patients only |
|  | 4 | SRP050190, SRP065041 (30, 61) | 383 | 1989-2014 | National (Canada) | Surveys from nationwide sites in Canada; enriched for CRO and AZM resistance |
|  | 5 | ERP010312 (27) | 714 | 2013 | International (Europe) | Survey from clinics and hospitals across 21 European countries |
|  | 6 | DRP004052 (28) | 204 | 2015 | National (Japan) | Survey from clinics in Kyoto and Osaka; male patients only |
|  | 7 | SRP111927 (63) | 398 | 2014-2015 | National (New Zealand) | Survey from nationwide diagnostic labs |
| *K. pneumoniae* | 8 | SRP102664 (14) | 1560 | 2011-2017 | Houston, TX (US) | Survey from citywide hospital system; enriched for β-lactam resistance |
| *A. baumannii* | 9 | SRP065910 (64) | 702 | 2000-2012 | National (US) | Survey from clinics and hospitals within the US military healthcare system |

CFX, cefixime; CRO, ceftriaxone; AZM, azithromycin

While it may be assumed that increased availability of paired genomic and phenotypic resistance data from a broader range of clinical populations will facilitate more accurate and reliable modeling (30), our results demonstrate that in predicting AZM resistance phenotypes for isolates from most datasets (with the exception of datasets 2 and 5), performance of classifiers trained on the aggregate dataset was not significantly better than performance of classifiers trained only on isolates from the dataset from which the test isolates were derived (*P <* 0.0001 and *P =* 0.002 for datasets 2 and 5, respectively, *P* = 0.008 for dataset 3, where the classifiers trained on the aggregate dataset had lower bACC than classifiers trained only on isolates from dataset 3, and *P >* 0.234 for all other datasets, **Fig 2a**). Further, there was substantial variation in performance of models trained on the aggregate dataset across testing sets, with models achieving significantly higher bACC for strains from datasets 3 and 4 than for strains from dataset 2 (*P <* 0.0009, **Fig 2a**), perhaps reflecting enrichment for AZM NS in these former datasets (**Table 1**). Additionally, with the exception of dataset 5, performance of AZM resistance classifiers trained only on isolates from the dataset from which the test isolates were derived was significantly higher than performance of classifiers trained on the aggregate dataset excluding isolates from the test dataset (*P* = 0.537 for dataset 5, *P <* 0.0005 for all other datasets, **Fig 2a**).

**Figure 2.**
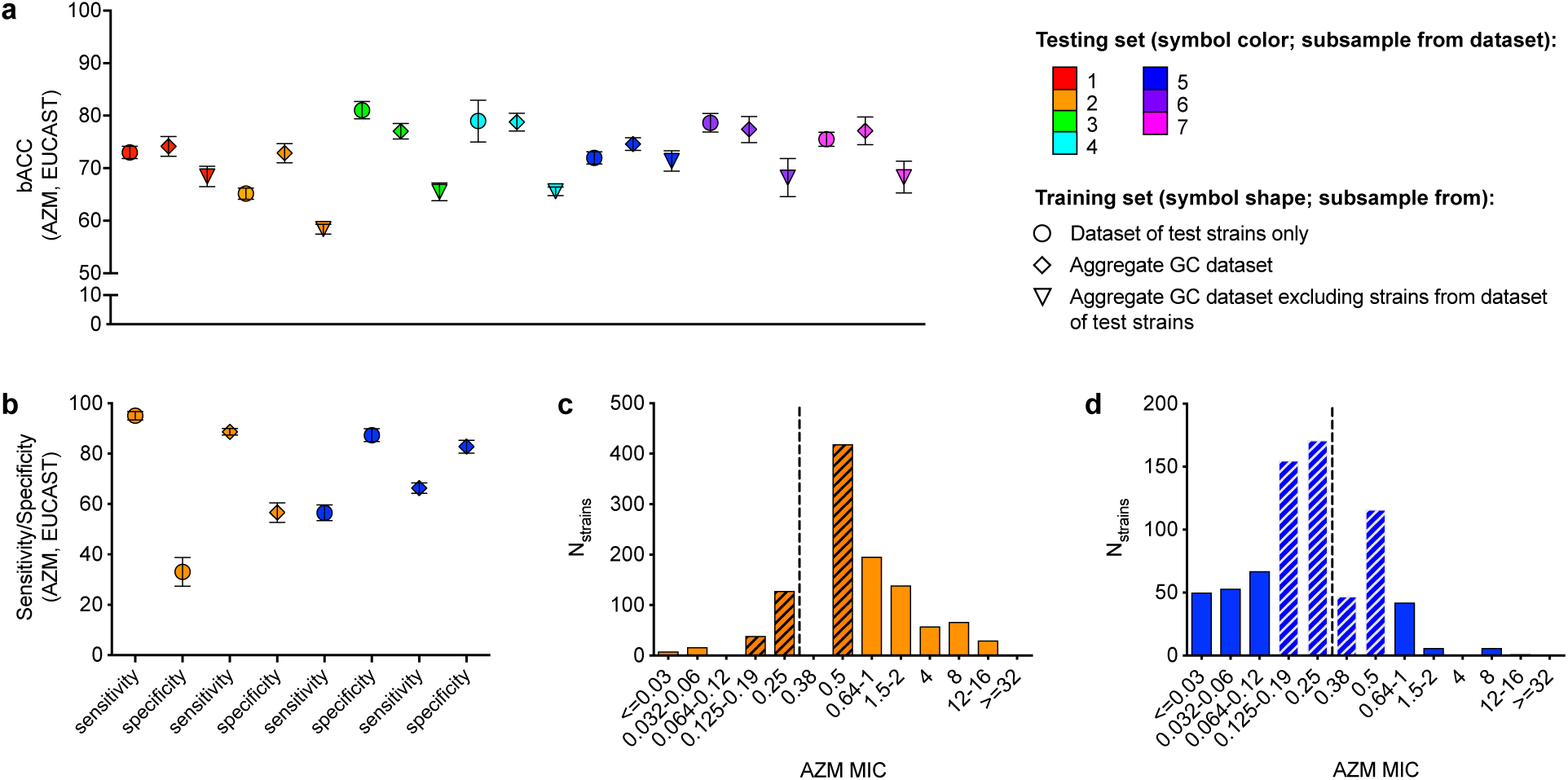
Differential performance of random forest classifiers across different datasets. **(a)** Mean balanced accuracy (bACC) with 95% confidence intervals of RF-C predictive models for gonococci (GC) azithromycin (AZM) non-susceptibility based on the EUCAST breakpoint. **(b)** Mean sensitivity and specificity with 95% confidence intervals of RF-C predictive models for GC AZM non-susceptibility in datasets 2 and 5. Histograms showing the distributions of AZM minimum inhibitory concentrations (MICs) in **(c)** dataset 2 and **(d)** dataset 5. Symbol colors in **(a)** and **(b)** indicate the dataset from which the testing set was derived, while symbol shape in **(a)** and **(b)** indicates the dataset from which the training set was derived. Hatching in **(c)** and **(d)** indicates MICs within one doubling dilution of the EUCAST breakpoint (designated by dashed lines).

Performance of RF classifiers trained and tested on dataset 2 was limited by low specificity, which was improved in models trained on the aggregate dataset (**Fig 2b**). The low specificity achieved by RF classifiers trained and tested on this dataset is likely due to the low representation of S strains, most of which were within one doubling dilution of the NS breakpoint (**Fig 2c**), and thus the more comprehensive representation of negative (S) data in the aggregate training set was associated with improved specificity. Conversely, performance of RF classifiers trained and tested on dataset 5 was more limited by low sensitivity, which was improved in models trained on the aggregate dataset (**Fig 2b**). This dataset had a low representation of strains with high AZM MICs (**Fig 2d**), and thus the more comprehensive representation of positive (NS) data in the aggregate training set was associated with improved sensitivity in predicting AZM NS for these strains. For both SCM and RF-C AZM resistance models across all datasets, there was a significant positive correlation between the ratio of model sensitivity to model specificity and the ratio of NS to S strains in the dataset (Pearson r > 0.98, *P <* 0.0001 [Pearson correlation] for both SCM and RF-C, **Fig S2a**).

On the other hand, while representation of strains with higher AZM MICs was also observed in other datasets (*i.e.*, datasets 1, 6, and 7) and was similarly reflected in the sensitivity-limited performance of RF classifiers trained and tested on these datasets (**Table S5**), AZM NS prediction accuracy for strains from these datasets was not improved by training classifiers on the aggregate dataset. Further, even after down-sampling two of the datasets with the most disparate MIC distributions, sample sizes, and model performance (datasets 2 and 4) such that the number of strains and AZM MIC distributions were identical between the two datasets (**Fig S2b**), there was still a significant difference in AZM NS prediction accuracy of models trained and tested on these different datasets (**Fig S2c,** *P* < 0.004). Together, these results demonstrate that resistance model performance may be strongly associated with the distributions of both resistance phenotypes and genetic features and thus can be highly population-specific.

### ML prediction models of antibiotic susceptibility / non-susceptibility outperform MIC models

Gonococcal CIP and AZM MICs were dichotomized by both EUCAST and CLSI breakpoints to assess the impact of variation in MIC breakpoints on model performance. As the EUCAST and CLSI breakpoints for CIP in gonococci are within a single doubling dilution and the vast majority of isolates have much lower or higher CIP MICs (**Fig 1a**), >99% of isolates in the aggregate dataset were consistently S or NS by both breakpoints. Of the 23 isolates with MICs between the two breakpoints, 18 had MICs derived from Etests of 0.032 μg/mL or 0.047 μg/mL, making their classification relative to the EUCAST breakpoint of 0.03 μg/mL ambiguous. In contrast, the EUCAST and CLSI breakpoints for AZM in gonococci are separated by two doubling dilutions, and for many isolates, the AZM MIC was within this range (**Fig 1b**). As such, only 67% of isolates in the aggregate dataset were consistently S or NS by both breakpoints. CIP NS classifier performance was either identical or nearly identical for both breakpoints in the aggregate and most individual gonococcal datasets (**Fig 3a**). In contrast, the bACC of AZM NS prediction by both SCM and RF classifiers based on the CLSI breakpoint was significantly higher than for those based on the EUCAST breakpoint across all gonococcal datasets assessed by both breakpoints (*P <* 0.0001, **Fig 3b**).

**Figure 3.**
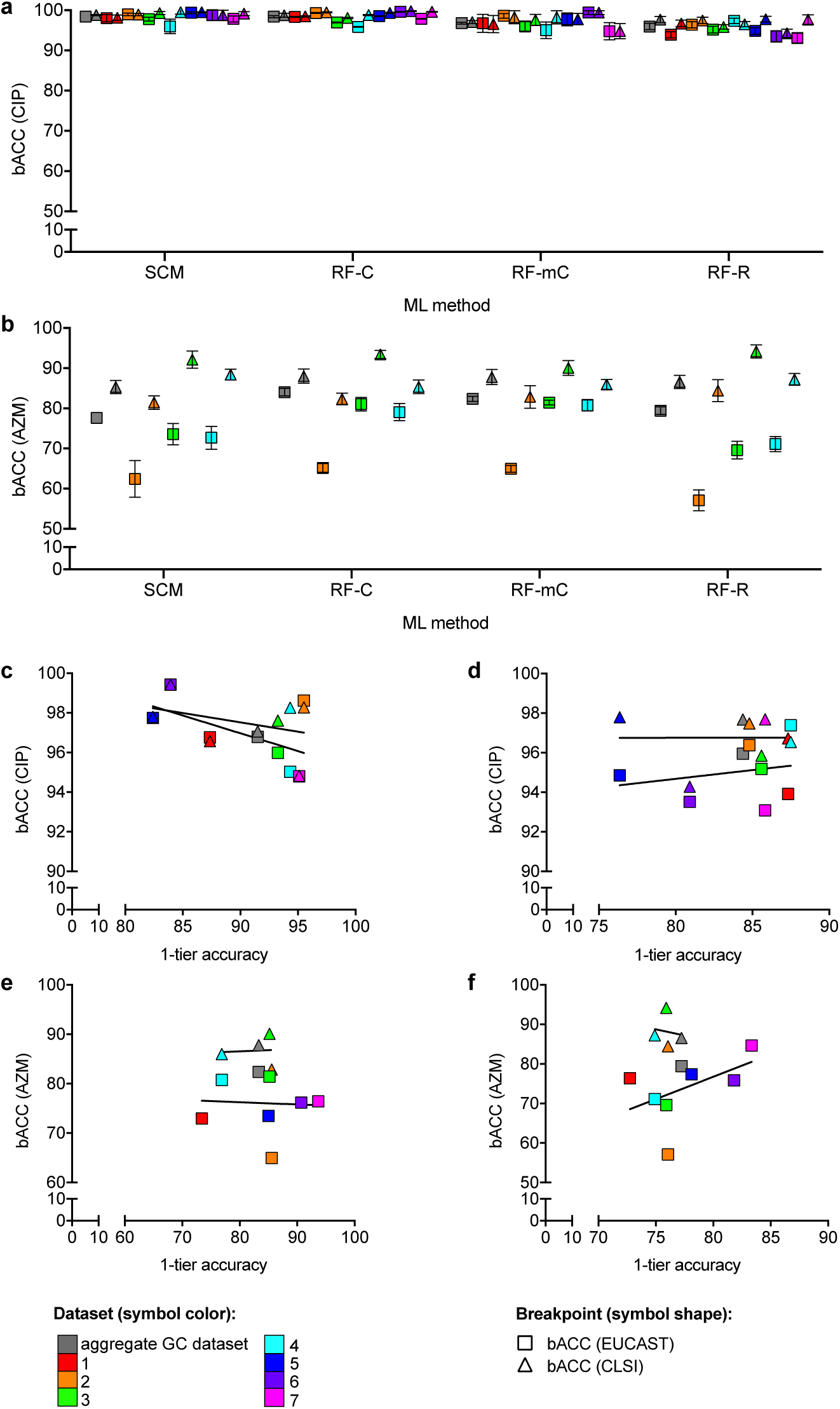
Differential performance of machine learning-based prediction models based on different resistance metrics in gonococci. Mean balanced accuracy (bACC) with 95% confidence intervals of predictive models for **(a)** ciprofloxacin non-susceptibility (CIP NS) across all datasets and **(b)** azithromycin (AZM) NS for all datasets for which both NS breakpoints were evaluated. Scatter plots comparing the mean 1-tier accuracy to the mean bACC for each gonococcal dataset derived from **(c-d)** CIP and **(e-f)** AZM minimum inhibitory concentration (MIC) prediction models by **(c,e)** random forest multi-class classification and **d,f** random forest regression. Symbol colors in **(a-f)** indicate the datasets from which the training and testing sets were derived. Symbol shapes in **(a-f)** indicate the NS breakpoint. The line of best fit for each of the breakpoints is indicated in **(c-f)**. SCM, set covering machine; RF-C, random forest binary classification; RF-mC, random forest multi-class classification; RF-R, random forest regression.

To assess the performance of MIC prediction models relative to binary S/NS resistance phenotype classifiers, RF-mC and RF-R models were trained and evaluated for CIP and AZM MIC prediction in gonococci. Average exact match rates between predicted and phenotypic MICs ranged from 64-86% and 54-78% by RF-mC and RF-R, respectively, for CIP, and from 24-60% and 45-65%, respectively, for AZM (**Tables S4-S5**). Average 1-tier accuracies (the percentage of isolates with predicted MICs within one doubling dilution of phenotypic MICs) were substantially higher but also varied widely across datasets and between the two MIC prediction methods (ranging from 82%-96% and 76-87% by RF-mC and RF-R, respectively, for CIP, and from 73-94% and 73-83%, respectively, for AZM; **Tables S4-S5**). There was no consistent or significant relationship across the different datasets between MIC prediction accuracy (exact match or 1-tier accuracy) and bACC for either drug by either MIC prediction method (**Fig 3c-f**). Further, for both drugs by both breakpoints in the aggregate gonococcal dataset, binary RF-C models had equivalent or significantly higher bACC than RF-mC and RF-R MIC prediction models (*P* > 0.175 for AZM NS by the CLSI breakpoint by RF-C compared to RF-mC or RF-R, P < 0.017 for all others, **Tables S4-S5**).

### Species with high genomic diversity pose challenges to ML-based antibiotic resistance prediction

Increasing genomic diversity, or an increasing ratio of genomic features (*e.g.*, *k*-mers) to observations (*e.g.*, genomes), may present an additional challenge for ML-based prediction of antibiotic resistance (12). To investigate ML-based antibiotic resistance prediction across species with different levels of genomic diversity, SCM and RF-C were used to model CIP NS in *K. pneumoniae* and *A. baumannii*, two species with genomic diversity (*i.e.*, ratio of unique 31-mers to number of genomes) several times that of gonococci (**Fig 4a-b**). SCM classifiers trained on and used to predict CIP NS for *K. pneumoniae* achieved significantly lower accuracy than all of the gonococcal datasets (*P* < 0.0001, **Fig 4c**), while SCM classifiers trained on and used to predict CIP NS for *A. baumannii* achieved significantly lower accuracy than gonococcal datasets 3-5 and 7 (*P* < 0.033) and roughly equivalent accuracy to gonococcal datasets 1-2 and 6, as well as the aggregate gonococcal dataset (*P >* 0.059, **Fig 4c**). The performance of RF-C models was significantly lower for both *K. pneumoniae* and *A. baumannii* compared to all gonococcal datasets (*P* < 0.0001, **Fig 4d**).

**Figure 4.**
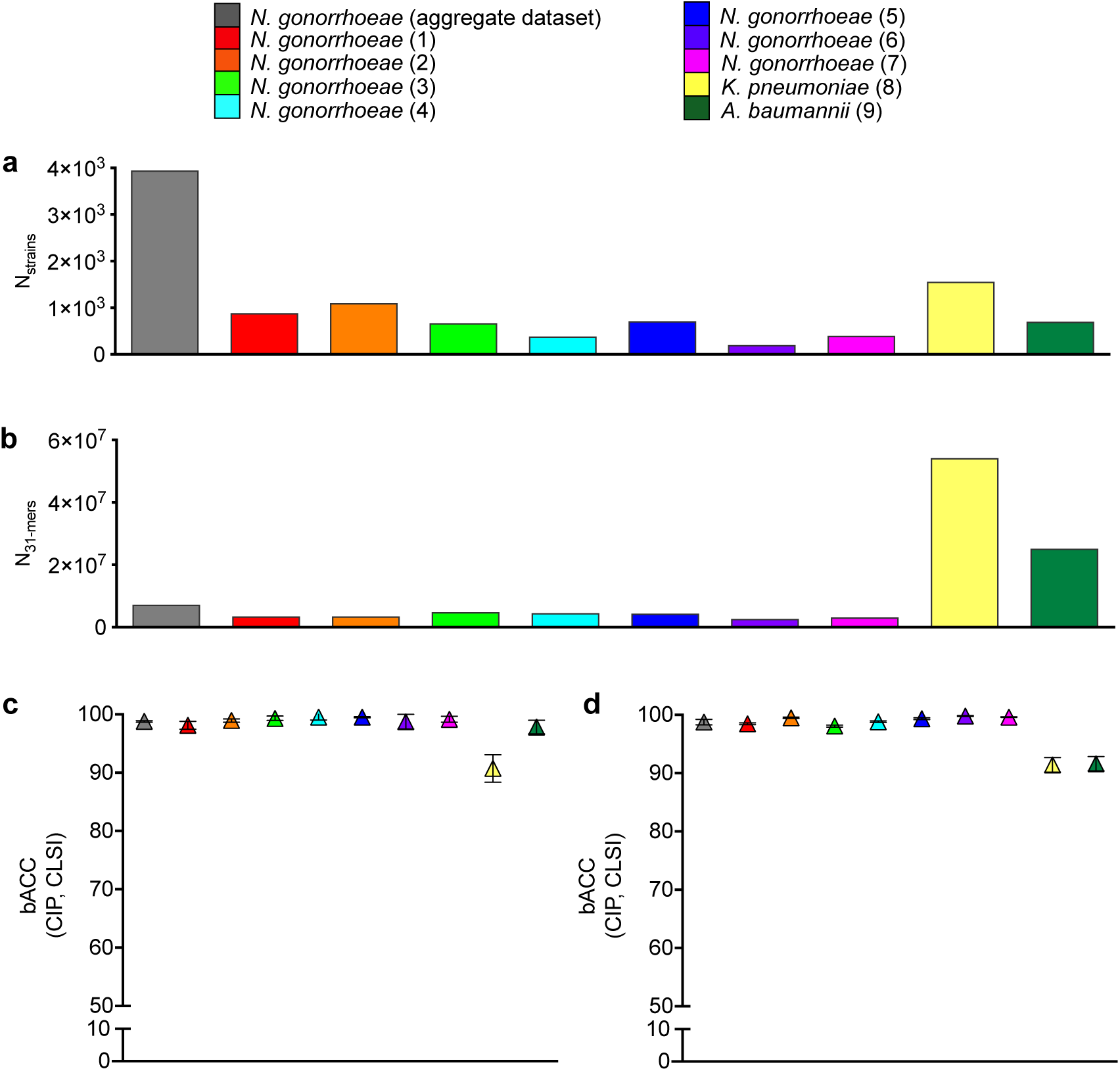
*K. pneumoniae* and *A. baumannii* datasets are associated with higher genetic diversity and lower performance of resistance prediction models. Number of **(a)** strains and **(b)** unique 31-mers present in the genomes of at least two strains in each dataset. Mean balanced accuracy (bACC) with 95% confidence intervals achieved by **c)** set covering machine and **d)** random forest classification models for ciprofloxacin (CIP) NS by the CLSI breakpoints across gonococcal, *K. pneumoniae,* and *A. baumannii* datasets.

While the SCM classifiers for CIP NS in *K. pneumoniae* performed significantly better on the training sets than the testing sets (**Table S4**, *P* < 0.0001), indicating that these models may be overfitted, there was no significant difference between RF-C model performance on training and testing sets for either *K. pneumoniae* or *A. baumannii* (*P* > 0.194), suggesting that overfitting alone cannot explain the variable classifier performance across different species. Down-sampling *K. pneumoniae* and *A. baumannii* to match the CIP MIC distributions of the gonococcal datasets was infeasible due to the narrow range of MICs tested for the former two species (**Table S7**). However, even after down-sampling to equalize the number of S and NS strains within each dataset (**Table S6, Fig. S3a-b**), performance of *K. pneumoniae* and *A. baumannii* CIP NS classifiers was still significantly lower than that of gonococcal CIP NS classifiers, with the exception of SCM classifiers based on the down-sampled *K. pneumoniae* dataset, which performed roughly equivalently to SCM classifiers based on gonococcal datasets 2 and 6 (P > 0.07 for the SCM classifiers based on the down-sampled *K. pneumoniae* dataset compared to SCM classifiers based on gonococcal datasets 2 and 6; P < 0.0004 for all other comparisons, **Figure S3c**).

Direct association based on GyrA codon 83 mutations (equivalent to codon 91 in gonococci) alone predicted CIP NS in *K. pneumoniae* with 86% sensitivity and 99% specificity, and thus had a marginally higher bACC (92.5%) than for the SCM classifiers and a substantially higher bACC than the RF classifiers. Similarly, for *A. baumannii*, GyrA codon 81 mutations (equivalent to codon 91 in gonococci) alone predicted CIP NS in with 97% sensitivity and 98% specificity, and thus with a roughly equivalent bACC (97.5%) to the SCM classifiers and a substantially higher bACC than the RF classifiers.

## Discussion

ML offers an opportunity to leverage WGS data to aid in development of rapid molecular diagnostics. While more comprehensive sampling of methods and parameters will be necessary to optimize model performance, we demonstrate that multiple factors beyond ML methods and parameters can affect model performance, reliability, and interpretability. Our results affirmed that drugs associated with complex and/or diverse resistance mechanisms present challenges to ML-based prediction of resistance phenotypes and that sampling frame (*i.e.*, temporal range, geographic range, and/or sampling approach) can substantially affect performance of such predictive models. We demonstrated significant variability in performance and potential clinical utility of predictive models based on different resistance metrics and further showed that the capacity to model antibiotic resistance may be highly variable across different species.

### Variable performance of ML-based resistance prediction models by antibiotic

Genotype-based resistance diagnostics have largely focused more on evaluating the presence of resistance determinants and less on predicting the susceptibility profile of a given isolate (8). However, in clinical settings where the empirical presumption is of resistance, prediction that an isolate is susceptible to an antibiotic may be more important in guiding treatment decisions. As such, the clinical utility of a genotype-based resistance diagnostic may be determined by its capacity to accurately predict susceptibility phenotype for multiple drugs.

While variable performance of ML-based predictive models has been observed across different drugs (7, 8, 10, 11, 14, 15), it has often been attributed to dataset size and/or imbalance (7, 14, 15). Further, while it is more difficult to predict resistance phenotypes from genotypes for drugs that are associated with unknown, multifactorial, and/or diverse resistance mechanisms than for drugs for which resistance can largely be attributed to a single variant (14, 29), this caveat has been presented specifically as a limitation of models based on known resistance loci in comparison to unbiased machine learning-based MIC prediction using genome-wide feature sets (14). However, by comparing performance of predictive models based on genome-wide feature sets between CIP and AZM across multiple gonococcal datasets, we showed that even with relatively large and phenotypically balanced datasets, ML algorithms cannot necessarily be expected to successfully model complex and/or diverse resistance mechanisms, particularly given that the representation of these resistance mechanisms in training datasets is *a priori* unknown.

As a high proportion of reported AZM MICs in gonococci are within 1-2 doubling dilutions of the NS breakpoints, it is possible that the inferior performance of AZM classifiers is partly attributable to errors and/or variations in MIC testing. However, given the noise of phenotypic MIC testing even with standardized protocols (32), this may be an inherent limitation of NS classifiers when low-level resistance is common. Further, while we show that removing strains with MICs ≤2 doubling dilutions from the breakpoints improved AZM classifier performance compared to AZM models trained and tested on the full dataset, performance of AZM classifiers trained and tested on this restricted dataset was still significantly lower than that of CIP classifiers, suggesting that additional drug-specific factors, such resistance mechanism diversity and/or complexity, can constrain classifier performance.

### Impact of demographic, geographic, and timeframe sampling bias on ML model predictions of antibiotic resistance

Sampling bias presents a substantial challenge in any predictive modeling, and sampling from limited patient demographics or during limited time periods may have considerable effects on the distributions of resistance phenotypes and resistance mechanisms (33, 34). For example, in TB, the RpoB I491F mutation that has been associated with failure of commercial RIF resistance diagnostic assays, including the GeneXpert MTB/RIF assay, reportedly accounted for <5% of TB RIF resistance in most countries, but, in Swaziland was found to be present in up to 30% of MDR-TB (35). Further, as the focus with statistical classifiers is building models from feature sets that can accurately predict an outcome, rather than understanding the association between each of the features and the outcome, potential confounding effects from factors such as population structure (36–38) or correlations among resistance profiles of different drugs (13) are rarely considered.

By comparing performance of AZM NS classifiers across multiple training and testing sets, we showed significant variation in performance of classifiers trained on a large and diverse global collection across testing sets from different sampling frames. In some cases of imbalanced datasets, models trained on datasets with a more comprehensive representation of resistance phenotypes improve prediction accuracy. Our results further demonstrate that the direction of dataset imbalance (*i.e.*, the ratio of NS to S strains) is significantly correlated with the direction of model performance (*i.e.*, the ratio of sensitivity to specificity), suggesting that, for example, optimizing sensitivity of predictive models for drugs with low prevalence of NS strains may require substantial enrichment of NS strains and/or down-sampling of S strains. However, while differential classifier performance among different datasets may be partially attributable to differential MIC distributions, our results also show variable classifier performance between datasets even in the case of identical MIC distributions (and sample size) and further suggest that heavier sampling across more geographic regions cannot necessarily be expected to significantly improve model performance, as models trained on the aggregate global gonococcal dataset did not improve prediction accuracy for most datasets.

This, together with decreased performance when excluding isolates from the dataset from which the isolates being tested were derived, suggests that factors such as population-specific resistance mechanisms, genetic divergence at resistance loci, and/or confounding effects may constrain model reliability across populations, particularly in the case of drugs like AZM with complex and/or diverse resistance mechanisms, where a substantial portion of the model may be overfit, or based on confounding factors or noise, rather than biologically-meaningful resistance variants. Further, it should be noted that MIC testing methods varied between some datasets (and between strains within dataset 5), and such variations may represent an additional confounding factor influencing classifier performance. Thus, both incorporation of methods to correct for potentially confounding factors, such as population structure, as have been introduced for genome-wide associate studies [15-17], and increased availability of paired WGS and antibiotic susceptibility data produced by consistent standardized protocols may improve reliability of machine learning-based prediction of antibiotic resistance across different populations.

### ML resistance prediction model performance varies by NS breakpoints and by categorical vs MIC-based resistance metrics

While measurement of MICs is vital for surveillance and investigation of resistance mechanisms, resistance breakpoints that relate *in vitro* MIC measurements to expected treatment outcomes inform clinical decision-making. However, standard breakpoints for NS to a given drug in a given species are often informed less by treatment outcome data, but rather factors such as pharmacokinetics and MIC distributions that can fail to account for a variety of intra-host conditions that could influence drug efficacy (39–42). Recent studies have shown that isolates that are classified as susceptible by standard breakpoints but have higher MICs are associated with a greater risk of treatment failure than isolates with lower MICs (43). Further, resistance breakpoints and testing protocols can vary across different organizations, and thus incongruence across phenotypic information included in the training data may introduce additional sources of error in predictive modeling. By comparing performance of predictive models of CIP and AZM NS based on EUCAST and CLSI breakpoints, we demonstrated breakpoint-specific performance of models. For CIP, such breakpoint-specific performance is likely largely attributable to variations in MIC testing protocols and thus ambiguous classification of some strains by the EUCAST breakpoint. On the other hand, the substantially lower performance of all AZM models based on the EUCAST breakpoint compared to those based on the CLSI breakpoint suggests that many isolates with AZM MICs between the two breakpoints lack genetic signatures that contribute to high model performance. While the clinical relevance of AZM MICs between these two breakpoints in gonococci is unclear, these isolates may be more likely to be associated with AZM treatment failure than isolates with lower MICs, and thus evaluation of classifiers using only higher breakpoints may misrepresent their diagnostic value, particularly in the absence of sufficient treatment outcome data.

Models that predict MICs provide more refined output than a binary classifier but generally achieve low rates of exact matches between phenotypic and predicted MICs and even fairly variable 1-tier accuracies (14, 15, 29). Given the noise in phenotypic MIC testing (32) and the potential lack of discriminating genetic features between isolates with MICs separated by 1-2 doubling dilutions (14), MIC prediction models may be unlikely to provide much better resolution than binary S/NS classifiers. Even if MIC predictions could provide additional resolution, the most important criterion of such a diagnostic would likely still be its ability to correctly predict resistance phenotypes relative to a clinically relevant breakpoint. Thus, performance of MIC prediction models with respect to breakpoints may be the biggest determinant of their diagnostic utility. By building MIC prediction models for CIP and AZM in gonococci, we observed low rates of exact matches between phenotypic and predicted MICs and variable 1-tier accuracies, with no relationship between 1-tier accuracy and categorical agreement (*i.e.*, prediction accuracy relative to NS breakpoints). Further, binary classifiers performed equivalently or better than MIC prediction models.

### ML antibiotic resistance prediction model success varies across species

Bacterial species with high genomic diversity (*e.g.*, open pangenomes) present additional challenges to ML-based prediction of antibiotic resistance. Increased resistance mechanism complexity and greater inter-isolate variation in resistance mechanisms require more intensive sampling to capture a significant portion of the resistome (47). On the technical side, even for heavily sampled species, when using whole genome feature sets, the number of genetic features (*e.g.*, k-mers or SNPs) will always be much larger than the number of observations (isolates), increasing the risk of overfitting (a situation that arises with so-called ‘fat data’; (12)). This raises concern in species with open pangenomes, as the ratio of genetic features to the number of genomes is larger and the number of unique genetic features per number of genomes does not plateau. By comparing classifier performance in predicting CIP NS across gonococci, *K. pneumoniae*, and *A. baumannii*, we show that classifiers generally did not perform as well for species with open genomes (*K. pneumoniae* or *A. baumannii*) as for gonococci. Further, while a single GyrA mutation could explain the majority of CIP NS across all species evaluated here, unlike in gonococci and *A. baumannii* where this mutation explained ≥97% of CIP NS, 14% of CIP NS in *K. pneumoniae* could not be explained by this mutation, suggesting increased CIP resistance mechanism diversity and/or complexity in this species. Increased sampling, different methods, and/or finer tuning of hyperparameters may yield increased prediction accuracy for drug resistance in species with open genomes. For example, Nguyen et al., 2018 reported a mean bACC of 98.5% (average VME and ME rates of 0.5% and 2.5%, respectively) using a decision tree-based extreme gradient boosting regression model to predict CIP MICs for the *K. pneumoniae* strains assessed here (14), and adjusting for confounding factors such as population structure or variation in MIC testing method may yield more consistent prediction accuracies across species. However, our results demonstrate clear variation in potential limitations of genotype-to-resistance-phenotype models across different species.

Given the biological and epidemiological disparities associated with resistance to different drugs in different clinical populations and bacterial species, and their evident impact on performance of predictive models, successful implementation of genotype-based resistance diagnostics will likely require sustained comprehensive sampling to ensure representation of complex, diverse, and/or novel resistance mechanisms, customized modeling, and incorporation of feedback mechanisms based on treatment outcome data. Further evaluation of additional ML methods and datasets may reveal more quantitative requirements and limitations associated with the application of genotype-to-resistance-phenotype predictive modeling in the clinical setting.

## Materials and Methods

### Isolate selection and dataset preparation

See **Table 1** for details of the datasets assessed and **Table S7** for per-strain information. All gonococcal datasets contained a minimum of 200 isolates with WGS (Illumina MiSeq, HiSeq, or NextSeq) and MICs available for both CIP and AZM (by agar dilution and/or Etest). Isolates lacking CIP and AZM MIC data were excluded. MIC testing methods varied within datasets, as reported (10-13, 17, 18, 29).

*K. pneumoniae* and *A. baumannii* datasets were selected based on the availability of isolates collected during a single survey that were tested for CIP susceptibility and whole genome sequenced using consistent platforms (in both cases, the BD-Phoenix system and either Illumina MiSeq or NextSeq).

MIC data were obtained from the associated publications, except in the cases of dataset 1 (NCBI Bioproject PRJEB10016; see **Table S7**) and dataset 9, which were obtained from the NCBI BioSample database (https://www.ncbi.nlm.nih.gov/biosample). Raw sequence data were downloaded from the NCBI Sequence Read Archive (https://www.ncbi.nlm.nih.gov/sra). Genomes were assembled using SPAdes (48) with default parameters, and assembly quality was assessed using QUAST (49). Contigs <200 bp in length and/or with <10x coverage were removed. Isolates with assembly N50s below two standard deviations of the dataset mean were removed.

### Evaluation of known resistance variants

Previously identified genetic loci associated with reduced susceptibility to CIP or AZM in gonococci are indicated in **Tables S1-S2**, respectively. The sequences of these loci were extracted from the gonococcus genome assemblies using BLAST (50) followed by MUSCLE alignment (51) to assess the presence or absence of known resistance variants. The presence or absence of quinolone resistance determining mutations in *gyrA* was similarly assessed in *K. pneumoniae* and *A. baumannii* assemblies. Presence or absence of gonococcal AZM resistance mutations in the multi-copy 23S rRNA gene was assessed using BWA-MEM(52) to map raw reads to a single 23S rRNA allele from the NCCP11945 reference isolate (NGK_rrna23s4), the Picard toolkit (http://broadinstitute.github.io/picard) to identify duplicate reads, and Pilon (53) to determine the mapping quality-weighted percentage of each nucleotide at the sites of interest.

### ML-based prediction of resistance phenotypes

Predictive modeling was carried out using SCM and RF algorithms, implemented in the Kover (11, 12) and ranger (54) packages, respectively. K-mer profiles (abundance profiles of all unique words of length k in each genome) were generated from the assembled contigs using the DSK k-mer counting software (55) with k=31, a length commonly used in bacterial genomic analysis (11, 12, 36, 56). For each dataset, 31-mer profiles for all strains were combined using the combinekmers tool implemented in SEER (36), removing 31-mers that were not present in more than one genome in the dataset. Final matrices used for model training and prediction were generated by converting the combined 31-mer counts for each dataset into presence/absence matrices. For each SCM binary classification analysis (using S/NS phenotypes based on the two different breakpoints for each drug), the best conjunctive and/or disjunctive model using a maximum of five rules was selected using five-fold cross-validation, testing the suggested broad range of values for the trade-off hyperparameter of 0.1, 0.178, 0.316, 0.562, 1.0, 1.778, 3.162, 5.623, 10.0, and 999999.0 to determine the optimal rule scoring function (http://aldro61.github.io/kover/doc_learning.html). In order to assess binary classification across multiple methods, RF was also used to build binary classifiers (RF-C) using S/NS phenotypes. Further, to compare performance of binary classifiers to MIC prediction models, RF was used to build multi-class classification (RF-mC) and regression (RF-R) models based on log_2_(MIC) data. For all RF analyses, forests were grown to 1000 trees using node impurity to assess variable importance and five-fold cross-validation to determine the most appropriate hyperparameters (yielding the highest bACC or 1-tier accuracy for NS- or MIC-based models, respectively), testing maximum tree depths of 5, 10, 100, and unlimited and mtry (number of features to split at each node) values of 1000, 10000, and either *√p* or *p*/3, for classification and regression models, respectively, where *p* is the total number of features (31-mers) in the dataset. While a grid search would enable assessment of more combinations of different hyperparameter values and thus finer tuning of hyperparameters, such an approach is computationally prohibitive on datasets of this size. To standardize reported MIC ranges across datasets, CIP MICs ≤0.008 μg/mL or ≥32 μg/mL were coded as 0.008 μg/mL or 32 μg/mL, respectively, and AZM MICs ≤0.008 μg/mL or ≥32 μg/mL were coded as 0.03 μg/mL or 32 μg/mL, respectively.

The set of SCM and RF analyses performed are indicated in **Tables S3** and **S6.** For each of the seven individual gonococcal datasets, as well as the aggregate gonococcal dataset (all gonococcal datasets combined, removing duplicate strains) and the *K. pneumoniae* and *A. baumannii* datasets, training sets consisted of random sub-samples of two-thirds of isolates from the dataset indicated (maintaining proportions of each resistance phenotype from the original dataset), while the remaining isolates were used to test performance of the model. Each set of analyses (for each combination of dataset/drug/resistance metric/ML algorithm) was performed on 10 replicates, each with a unique randomly partitioned training and testing set. For all gonococcal datasets, separate models were trained and tested using the EUCAST (57) and CLSI (58) breakpoints for NS to CIP. Four of the *N. gonorrhoeae* datasets had insufficient (<15) NS isolates by the CLSI breakpoint for AZM non-susceptibility and thus were only assessed at the EUCAST AZM breakpoint. CIP MICs for the *K. pneumoniae* isolates were not available in the range of the EUCAST breakpoint (0.25 μg/mL), and thus only the CLSI breakpoint for NS (>1 μg/mL) was assessed. For *A. baumannii*, the EUCAST and CLSI breakpoints for ciprofloxacin NS are the same (>1 μg/mL). Due to the very limited range of MICs within the BD-Phoenix testing thresholds and thus the CIP MICs available for *K. pneumoniae* and *A. baumannii*, predictive models based on MICs were not generated for these species. For analyses in **Table S6** where datasets were down-sampled to equalize MIC distributions between datasets or the number of S and NS strains within datasets, the required number of strains from the over-represented class(es) were selected at random for removal.

Model performance was assessed by sensitivity (1 – VME rate), specificity (1 – ME rate), and aggregate bACC (the average of the sensitivity and specificity (59)). bACC was used as an aggregate measure of model performance as, unlike metrics such as raw accuracy, error rate, and F1 score, it provides a balanced representation of false positive and false negative rates, even in the case of dataset imbalance. For MIC prediction models, the percentage of isolates with predicted MICs exactly matching the phenotypic MICs (rounding to the nearest doubling dilution, in the case of regression models), as well as the percentage of isolates with predicted MICs within one doubling dilution of phenotypic MICs (1-tier accuracy), were also assessed. In order to account for variations in MIC testing methods and thus in the dilutions assessed, criteria for exact match rates and 1-tier accuracies were relaxed to include predictions within 0.5 doubling dilutions or 1.5 doubling dilutions, respectively, of the phenotypic MIC. Mean and 95% confidence intervals for all metrics were calculated across the 10 replicates for each analysis. Differential model performance between datasets or methods was evaluated by comparing mean bACC between sets of replicates by two-tailed unpaired t-tests with Welch’s correction for unequal variance (α=0.05). Unless otherwise noted, all *P*-values are derived from these unpaired t-tests. Relationships between MIC prediction accuracy and bACC and between dataset imbalance and model performance were assessed by Pearson correlation (α=0.05).

## Acknowledgements

We thank Jung-Eun Shin, Mark Labrador, and members of the Grad Lab for helpful discussion, and Julie Schillinger and Preeti Pathela for assistance identifying, selecting, and characterizing the isolates from New York City. The authors declare no competing interests.

## Supplementary Tables and Figures

**Table S1.** Genetic variants previously associated with ciprofloxacin resistance in *N. gonorrhoeae*.

**Table S2.** Genetic variants previously associated with azithromycin resistance in *N. gonorrhoeae*.

**Table S3.** Summary of approach in the primary set covering machine and random forest analyses.

**Table S4.** Performance (mean with 95% confidence intervals) of predictive models for ciprofloxacin resistance from the primary set covering machine and random forest analyses.

**Table S5.** Performance (mean with 95% confidence intervals) of predictive models for azithromycin resistance from the primary set covering machine and random forest analyses.

**Table S6.** Summary of approach in the additional random forest analyses for assessment of sampling bias.

**Table S7.** Study ID, machine learning dataset(s), antibiotic susceptibility testing (AST) methods, azithromycin (AZM) and ciprofloxacin (CIP) minimum inhibitory concentrations (MICs) for all strains assessed.

**Figure S1. MIC distribution influences classifier results but cannot explain all drug-specific classifier performance.** Histograms showing azithromycin (AZM) minimum inhibitory concentration (MIC) distributions for the aggregate gonococcal dataset after down-sampling to remove all strains with MICs ≤2 doubling dilutions of the **(a)** EUCAST or **(b)** CLSI breakpoint. **(c)** Mean balanced accuracy (bACC) with 95% confidence intervals of SCM RF-C predictive models trained and tested on down-sampled aggregate gonococcal datasets.

**Figure S2. Dataset imbalance influences classifier results but cannot explain all dataset-specific classifier performance. (a)** Scatter plot showing the relationship between the ratio of azithromycin (AZM) non-susceptible (NS) strains to susceptible (S) strains (by the EUCAST breakpoint) in each dataset and the ratio of sensitivity to specificity achieved by set covering machine (SCM) and random forest binary classification (RF-C) methods. **(b)** Histogram showing the AZM minimum inhibitory concentration (MIC) distribution for both datasets 2 and 4 after down-sampling to equalize number of strains and MIC distributions between datasets. **(c)** Mean balanced accuracy (bACC) with 95% confidence intervals of RF-C predictive AZM NS models trained and tested on down-sampled datasets 2 and 4. Symbol colors in **(a)** indicated the machine learning (ML) method. Symbol colors **(b)** indicate the down-sampled dataset from which the training and testing sets were derived.

**Figure S3. Down-sampling to balance resistance phenotypes does ameliorate cross-species variation in classifier performance.** Number of **(a)** strains and **(b)** unique 31-mers present in the genomes of at least two strains in each dataset, after down-sampling the *K. pneumoniae* and *A. baumannii* datasets to equalize the number of S and NS strains within each dataset. Mean balanced accuracy (bACC) with 95% confidence intervals achieved by **c)** set covering machine and **d)** random forest classification models for ciprofloxacin (CIP) NS by the CLSI breakpoints across gonococcal, down-sampled *K. pneumoniae,* and down-sampled *A. baumannii* datasets.

