## Supplemental Table 1 for "Evaluation of parameters affecting performance and reliability of machine learning-based antibiotic susceptibility testing from whole genome sequencing data"

**Table S1.** Genetic variants previously associated with ciprofloxacin resistance in *N. gonorrhoeae.*

|  | **Locus** | *gyrA* | | *parC* | | | | *parE* |
| --- | --- | --- | --- | --- | --- | --- | --- | --- |
|  | **NS marker** | S91F | D95A/G/N | D86N | S87I/R/W | S88P | E91G/K/Q | G410V  (1) |
|  | **Reference** | (2, 3) | (2-4) | (2, 3) | (2-4) | (2, 3) | (2-5) |  |
| **Aggregate GC dataset** | S (EUCAST) | 16 (0.8%) | 17 (0.8%) | 0 (0%) | 16 (0.8%) | 0 (0%) | 4 (0.2%) | 0 (0%) |
|  | NS (EUCAST) | 1779 (98%) | 1769 (97.5%) | 102 (5.6%) | 1386 (76.4%) | 55 (3%) | 254 (14%) | 2 (0.1%) |
|  | S (CLSI) | 21 (1%) | 21 (1%) | 0 (0%) | 19 (0.9%) | 0 (0%) | 4 (0.2%) | 0 (0%) |
|  | NS (CLSI) | 1774 (99%) | 1765 (98.5%) | 102 (5.7%) | 1383 (77.2%) | 55 (3.1%) | 254 (14.2%) | 2 (0.1%) |
| **1** | S (EUCAST) | 7 (1.1%) | 7 (1.1%) | 0 (0%) | 6 (0.9%) | 0 (0%) | 4 (0.6%) | 0 (0%) |
|  | NS (EUCAST) | 220 (96.9%) | 220 (96.9%) | 6 (2.6%) | 145 (63.9%) | 0 (0%) | 70 (30.8%) | 0 (0%) |
|  | S (CLSI) | 10 (1.5%) | 10 (1.5%) | 0 (0%) | 9 (1.4%) | 0 (0%) | 4 (0.6%) | 0 (0%) |
|  | NS (CLSI) | 217 (97.7%) | 217 (97.7%) | 6 (2.7%) | 142 (64%) | 0 (0%) | 70 (31.5%) | 0 (0%) |
| **2** | S (EUCAST) | 4 (0.8%) | 4 (0.8%) | 0 (0%) | 3 (0.6%) | 0 (0%) | 0 (0%) | 0 (0%) |
|  | NS (EUCAST) | 594 (98.7%) | 591 (98.2%) | 7 (1.2%) | 548 (91%) | 1 (0.2%) | 29 (4.8%) | 0 (0%) |
|  | S (CLSI) | 6 (1.2%) | 4 (0.8%) | 0 (0%) | 3 (0.6%) | 0 (0%) | 0 (0%) | 0 (0%) |
|  | NS (CLSI) | 592 (98.8%) | 591 (98.7%) | 7 (1.2%) | 548 (91.5%) | 1 (0.2%) | 29 (4.8%) | 0 (0%) |
| **3** | S (EUCAST) | 3 (1%) | 3 (1%) | 0 (0%) | 3 (1%) | 0 (0%) | 0 (0%) | 0 (0%) |
|  | NS (EUCAST) | 367 (96.8%) | 364 (96.1%) | 18 (4.7%) | 319 (84.2%) | 2 (0.5%) | 20 (5.3%) | 2 (0.5%) |
|  | S (CLSI) | 3 (1%) | 3 (1%) | 0 (0%) | 3 (1%) | 0 (0%) | 0 (0%) | 0 (0%) |
|  | NS (CLSI) | 367 (99.7%) | 364 (98.9%) | 18 (4.9%) | 319 (86.7%) | 2 (0.5%) | 20 (5.4%) | 2 (0.5%) |
| **4** | S (EUCAST) | 0 (0%) | 0 (0%) | 0 (0%) | 0 (0%) | 0 (0%) | 0 (0%) | 0 (0%) |
|  | NS (EUCAST) | 190 (94.1%) | 182 (90.1%) | 10 (5.0%) | 160 (79.2%) | 4 (2%) | 6 (3.0%) | 1 (0.5%) |
|  | S (CLSI) | 0 (0%) | 0 (0%) | 0 (0%) | 0 (0%) | 0 (0%) | 0 (0%) | 0 (0%) |
|  | NS (CLSI) | 190 (98.4%) | 182 (94.3%) | 10 (5.2%) | 160 (82.9%) | 4 (2.1%) | 6 (3.1%) | 1 (0.5%) |
| **5** | S (EUCAST) | 2 (0.5%) | 3 (0.8%) | 0 (0%) | 4 (1.1%) | 0 (0%) | 0 (0%) | 0 (0%) |
|  | NS (EUCAST) | 339 (99.1%) | 340 (99.4%) | 31 (9.1%) | 230 (67.3%) | 5 (1.5%) | 78 (22.8%) | 0 (0%) |
|  | S (CLSI) | 2 (0.5%) | 3 (0.8%) | 0 (0%) | 4 (1.1%) | 0 (0%) | 0 (0%) | 0 (0%) |
|  | NS (CLSI) | 339 (99.7%) | 340 (100%) | 31 (9.1%) | 230 (67.6%) | 5 (1.5%) | 78 (22.9%) | 0 (0%) |
| **6** | S (EUCAST) | 0 (0%) | 0 (0%) | 0 (0%) | 0 (0%) | 0 (0%) | 0 (0%) | 0 (0%) |
|  | NS (EUCAST) | 172 (98.9%) | 172 (98.9%) | 24 (13.8%) | 145 (83.3%) | 45 (25.9%) | 5 (2.9%) | 0 (0%) |
|  | S (CLSI) | 0 (0%) | 0 (0%) | 0 (0%) | 0 (0%) | 0 (0%) | 0 (0%) | 0 (0%) |
|  | NS (CLSI) | 172 (98.9%) | 172 (98.9%) | 24 (13.8%) | 145 (83.3%) | 45 (25.9%) | 5 (2.9%) | 0 (0%) |
| **7** | S (EUCAST) | 2 (0.7%) | 2 (0.7%) | 0 (0%) | 2 (0.7%) | 0 (0%) | 0 (0%) | 0 (0%) |
|  | NS (EUCAST) | 124 (97.6%) | 125 (98.4%) | 10 (7.9%) | 48 (37.8%) | 0 (0%) | 53 (41.7%) | 0 (0%) |
|  | S (CLSI) | 2 (0.7%) | 3 (1.1%) | 0 (0%) | 2 (0.7%) | 0 (0%) | 0 (0%) | 0 (0%) |
|  | NS (CLSI) | 124 (98.4%) | 124 (98.4%) | 10 (7.9%) | 48 (38.1%) | 0 (0%) | 53 (42.1%) | 0 (0%) |

NS, non-susceptible; S, susceptible

*norM* T-35C promoter mutation (6) not present in any strains
