## Supplemental Table 2 for "Evaluation of parameters affecting performance and reliability of machine learning-based antibiotic susceptibility testing from whole genome sequencing data"

**Table S2.** Genetic variants previously associated with azithromycin resistance in *N. gonorrhoeae*.

|  | **Locus** | 23S rRNA | | | | | | *mtrR* | | | | | | | *ermB* | *ermC* | *mefAE* |
| --- | --- | --- | --- | --- | --- | --- | --- | --- | --- | --- | --- | --- | --- | --- | --- | --- | --- |
|  | **NS marker** | C2611T (1/4) | C2611T (2/4) | C2611T (3/4) | C2611T (4/4) | C2611T (any) | A2059G | "mtr120" | A-38C (mosaic promoter) | A-35 deletion | -10 TT insertion | A39T | G45D | truncation | presence | presence | presence |
|  | **Reference** | (1) | (1) | (1) | (1) | (1) | (2) | (3) | (4) | (5) | (6) | (7) | (7) | (8) | (9) | (9) | (6) |
| **Aggregate GC dataset** | S (EUCAST) | 6 (0.3%) | 0 (0%) | 0 (0%) | 0 (0%) | 6 (0.3%) | 0 (0%) | 4 (0.2%) | 9 (0.4%) | 677 (31.9%) | 18 (0.9%) | 663 (31.3%) | 339 (16%) | 91 (4.3%) | 0 (0%) | 0 (0%) | 1 (0.1%) |
|  | NS (EUCAST) | 11 (0.6%) | 21 (1.2%) | 35 (1.9%) | 264 (14.4%) | 331 (18.1%) | 7 (0.4%) | 5 (0.3%) | 51 (2.8%) | 1108 (60.7%) | 2 (0.1%) | 361 (19.8%) | 233 (12.8%) | 66 (3.6%) | 2 (0.1%) | 4 (0.2%) | 1 (0.1%) |
|  | S (CLSI) | 16 (0.5%) | 3 (0.1%) | 1 (0%) | 0 (0%) | 20 (0.6%) | 0 (0%) | 8 (0.2%) | 14 (0.4%) | 1545 (45.3%) | 20 (0.6%) | 934 (27.4%) | 460 (13.5%) | 140 (4.1%) | 0 (0%) | 2 (0.1%) | 2 (0.1%) |
|  | NS (CLSI) | 1 (0.2%) | 18 (3.4%) | 34 (6.3%) | 264 (49.3%) | 317 (59.1%) | 7 (1.3%) | 1 (0.2%) | 46 (8.6%) | 240 (44.8%) | 0 (0%) | 90 (16.8%) | 112 (20.9%) | 17 (3.2%) | 2 (0.4%) | 2 (0.4%) | 0 (0%) |
| **1** | S (EUCAST) | 2 (0.3%) | 0 (0%) | 0 (0%) | 0 (0%) | 2 (0.3%) | 0 (0%) | 0 (0%) | 2 (0.3%) | 162 (24.7%) | 1 (0.2%) | 176 (26.8%) | 152 (23.2%) | 28 (4.3%) | 0 (0%) | 0 (0%) | 0 (0%) |
|  | NS (EUCAST) | 0 (0%) | 0 (0%) | 0 (0%) | 3 (1.3%) | 3 (1.3%) | 0 (0%) | 0 (0%) | 5 (2.2%) | 94 (40.9%) | 0 (0%) | 104 (45.2%) | 15 (6.5%) | 2 (0.9%) | 0 (0%) | 0 (0%) | 0 (0%) |
|  | S (CLSI) | 2 (0.2%) | 0 (0%) | 0 (0%) | 0 (0%) | 2 (0.2%) | 0 (0%) | 0 (0%) | 3 (0.3%) | 250 (28.5%) | 1 (0.1%) | 280 (32%) | 167 (19.1%) | 30 (3.4%) | 0 (0%) | 0 (0%) | 0 (0%) |
|  | NS (CLSI)^1^ | 0 (0%) | 0 (0%) | 0 (0%) | 3 (30%) | 3 (30%) | 0 (0%) | 0 (0%) | 4 (40%) | 6 (60%) | 0 (0%) | 0 (0%) | 0 (0%) | 0 (0%) | 0 (0%) | 0 (0%) | 0 (0%) |
| **2** | S (EUCAST) | 0 (0%) | 0 (0%) | 0 (0%) | 0 (0%) | 0 (0%) | 0 (0%) | 0 (0%) | 0 (0%) | 101 (52.3%) | 0 (0%) | 45 (23.3%) | 24 (13%) | 27 (14%) | 0 (0%) | 0 (0%) | 0 (0%) |
|  | NS (EUCAST) | 8 (0.9%) | 10 (1.1%) | 23 (2.5%) | 131 (14.4%) | 172 (18.9%) | 2 (0.2%) | 0 (0%) | 28 (3.1%) | 559 (61.5%) | 2 (0.2%) | 103 (11.3%) | 167 (18.3%) | 8 (0.9%) | 0 (0%) | 0 (0%) | 0 (0%) |
|  | S (CLSI) | 8 (1%) | 2 (0.3%) | 0 (0%) | 0 (0%) | 10 (1.2%) | 0 (0%) | 0 (0%) | 2 (0.3%) | 553 (68.5%) | 2 (0.3%) | 100 (12.4%) | 102 (12.7%) | 33 (4.1%) | 0 (0%) | 0 (0%) | 0 (0%) |
|  | NS (CLSI) | 0 (0%) | 8 (2.7%) | 23 (7.8%) | 131 (44.4%) | 162 (54.9%) | 2 (0.7%) | 0 (0%) | 26 (8.8%) | 107 (36.3%) | 0 (0%) | 48 (16.3%) | 89 (30.2%) | 2 (0.7%) | 0 (0%) | 0 (0%) | 0 (0%) |
| **3** | S (EUCAST) | 2 (0.9%) | 0 (0%) | 0 (0%) | 0 (0%) | 2 (0.9%) | 0 (0%) | 0 (0%) | 0 (0%) | 95 (40.8%) | 1 (0.4%) | 65 (27.9%) | 38 (16.7%) | 15 (6.4%) | 0 (0%) | 0 (0%) | 0 (0%) |
|  | NS (EUCAST) | 1 (0.2%) | 11 (2.5%) | 12 (2.7%) | 133 (30.2%) | 157 (35.7%) | 4 (0.9%) | 3 (0.7%) | 22 (5%) | 290 (66.1%) | 1 (0.2%) | 59 (13.4%) | 47 (10.7%) | 23 (5.2%) | 2 (0.5%) | 2 (0.5%) | 0 (0%) |
|  | S (CLSI) | 2 (0.5%) | 0 (0%) | 1 (0.2%) | 0 (0%) | 3 (0.7%) | 0 (0%) | 2 (0.5%) | 2 (0.5%) | 269 (61.1%) | 2 (0.5%) | 76 (17.2%) | 61 (14%) | 17 (3.9%) | 0 (0%) | 0 (0%) | 0 (0%) |
|  | NS (CLSI) | 1 (0.4%) | 11 (4.8%) | 11 (4.8%) | 133 (57.6%) | 156 (67.5%) | 4 (1.7%) | 1 (0.4%) | 20 (8.7%) | 116 (50.2%) | 0 (0%) | 48 (20.8%) | 24 (10.4%) | 21 (9.1%) | 2 (0.9%) | 2 (0.9%) | 0 (0%) |
| **4** | S (EUCAST) | 2 (1.9%) | 0 (0%) | 0 (0%) | 0 (0%) | 2 (1.9%) | 0 (0%) | 4 (3.9%) | 0 (0%) | 43 (41.8%) | 0 (0%) | 24 (23.3%) | 18 (17.5%) | 2 (1.9%) | 0 (0%) | 0 (0%) | 0 (0%) |
|  | NS (EUCAST) | 1 (0.4%) | 10 (3.6%) | 10 (3.6%) | 116 (41.4%) | 127 (45.4%) | 4 (1.4%) | 1 (0.4%) | 17 (6.1%) | 177 (63.2%) | 0 (0%) | 44 (15.7%) | 25 (8.9%) | 22 (7.9%) | 2 (0.7%) | 3 (1.1%) | 0 (0%) |
|  | S (CLSI) | 2 (1.1%) | 0 (0%) | 1 (0.6%) | 0 (0%) | 3 (1.7%) | 0 (0%) | 4 (2.3%) | 1 (0.6%) | 108 (61.7%) | 0 (0%) | 29 (16.6%) | 24 (13.7%) | 4 (2.3%) | 0 (0%) | 1 (0.6%) | 0 (0%) |
|  | NS (CLSI) | 1 (0.5%) | 10 (4.8%) | 9 (4.3%) | 116 (55.8%) | 126 (60.6%) | 4 (1.9%) | 1 (0.5%) | 16 (7.7%) | 112 (53.9%) | 0 (0%) | 39 (18.8%) | 19 (9.1%) | 20 (9.6%) | 2 (1%) | 2 (1%) | 0 (0%) |
| **5** | S (EUCAST) | 1 (0.2%) | 0 (0%) | 0 (0%) | 0 (0%) | 1 (0.2%) | 0 (0%) | 1 (0.2%) | 3 (0.6%) | 170 (34.3%) | 16 (3.2%) | 179 (36.1%) | 34 (6.9%) | 36 (7.3%) | 0 (0%) | 0 (0%) | 1 (0.2%) |
|  | NS (EUCAST) | 1 (0.5%) | 0 (0%) | 2 (0.9%) | 6 (2.8%) | 9 (4.1%) | 0 (0%) | 0 (0%) | 0 (0%) | 143 (65.6%) | 0 (0%) | 76 (34.9%) | 7 (3.2%) | 6 (2.8%) | 0 (0%) | 1 (0.5%) | 1 (0.5%) |
|  | S (CLSI) | 2 (0.3%) | 0 (0%) | 0 (0%) | 0 (0%) | 2 (0.3%) | 0 (0%) | 1 (0.1%) | 3 (0.4%) | 303 (43.2%) | 16 (2.3%) | 252 (36%) | 41 (5.9%) | 42 (6%) | 0 (0%) | 1 (0.1%) | 2 (0.3%) |
|  | NS (CLSI)^1^ | 0 (0%) | 0 (0%) | 2 (15.4%) | 6 (46.2%) | 8 (61.5%) | 0 (0%) | 0 (0%) | 0 (0%) | 10 (76.9%) | 0 (0%) | 3 (23.1%) | 0 (0%) | 0 (0%) | 0 (0%) | 0 (0%) | 0 (0%) |
| **6** | S (EUCAST) | 0 (0%) | 0 (0%) | 0 (0%) | 0 (0%) | 0 (0%) | 0 (0%) | 0 (0%) | 1 (0.8%) | 50 (38.2%) | 0 (0%) | 11 (8.4%) | 39 (29.8%) | 0 (0%) | 0 (0%) | 0 (0%) | 0 (0%) |
|  | NS (EUCAST) | 1 (1.4%) | 1 (1.4%) | 0 (0%) | 2 (2.7%) | 4 (5.5%) | 1 (1.4%) | 2 (2.7%) | 0 (0%) | 56 (76.7%) | 0 (0%) | 8 (11%) | 9 (12.3%) | 0 (0%) | 0 (0%) | 0 (0%) | 0 (0%) |
|  | S (CLSI) | 1 (0.5%) | 1 (0.5%) | 0 (0%) | 0 (0%) | 2 (1%) | 0 (0%) | 2 (1%) | 1 (0.5%) | 104 (51.7%) | 0 (0%) | 19 (9.5%) | 47 (23.4%) | 0 (0%) | 0 (0%) | 0 (0%) | 0 (0%) |
|  | NS (CLSI)^1^ | 0 (0%) | 0 (0%) | 0 (0%) | 2 (66.7%) | 2 (66.7%) | 1 (33.3%) | 0 (0%) | 0 (0%) | 2 (66.7%) | 0 (0%) | 0 (0%) | 1 (33.3%) | 0 (0%) | 0 (0%) | 0 (0%) | 0 (0%) |
| **7** | S (EUCAST) | 0 (0%) | 0 (0%) | 0 (0%) | 0 (0%) | 0 (0%) | 0 (0%) | 1 (0.3%) | 3 (0.9%) | 82 (23.1%) | 0 (0%) | 175 (49.3%) | 41 (11.6%) | 14 (3.9%) | 0 (0%) | 0 (0%) | 0 (0%) |
|  | NS (EUCAST) | 0 (0%) | 0 (0%) | 0 (0%) | 2 (4.7%) | 2 (4.7%) | 0 (0%) | 0 (0%) | 0 (0%) | 22 (51.2%) | 0 (0%) | 20 (46.5%) | 1 (2.3%) | 1 (2.3%) | 0 (0%) | 0 (0%) | 0 (0%) |
|  | S (CLSI) | 0 (0%) | 0 (0%) | 0 (0%) | 0 (0%) | 0 (0%) | 0 (0%) | 1 (0.3%) | 3 (0.8%) | 102 (25.8%) | 0 (0%) | 195 (49.2%) | 42 (10.6%) | 15 (3.8%) | 0 (0%) | 0 (0%) | 0 (0%) |
|  | NS (CLSI)^1^ | 0 (0%) | 0 (0%) | 0 (0%) | 2 (100%) | 2 (100%) | 0 (0%) | 0 (0%) | 0 (0%) | 2 (100%) | 0 (0%) | 0 (0%) | 0 (0%) | 0 (0%) | 0 (0%) | 0 (0%) | 0 (0%) |

NS, non-susceptible; S, susceptible

*macAB* G-48-T promoter mutation (10), *ermA* (9), *ermF* (9), and *ereAB* (11) not present in any strains

^1^Azithromycin NS by the CLSI breakpoint was not assessed for dataset 1 or 5-7 due to low representation (<15) of NS strains.
