## Supplemental Table 3 for "Evaluation of parameters affecting performance and reliability of machine learning-based antibiotic susceptibility testing from whole genome sequencing data"

**Table S3.** Summary of approach in the primary set covering machine and random forest analyses.

| Species | Dataset^1^ | Drug | Resistance metric | ML algorithm |
| --- | --- | --- | --- | --- |
| *N. gonorrhoeae* | 1-7  Aggregate GC dataset | CIP | NS (EUCAST, >0.03 μg/mL) | SCM, RF-C |
|  |  |  | NS (CLSI, >0.06 μg/mL) | SCM, RF-C |
|  |  |  | log_2_(MIC) | RF-mC, RF-R |
|  |  | AZM | NS (EUCAST, >0.25 μg/mL) | SCM, RF-C |
|  |  |  | log_2_(MIC) | RF-mC, RF-R |
|  | 2-4  Aggregate GC dataset | AZM | NS (CLSI, >1 μg/mL) | SCM, RF-C |
| *K. pneumoniae* | 8 | CIP | NS (CLSI, >1 μg/mL) | SCM, RF-C |
| *A. baumannii* | 9 | CIP | NS (EUCAST/CLSI, >1 μg/mL) | SCM, RF-C |

GC, gonococcal; CIP, ciprofloxacin; AZM, azithromycin; NS, non-susceptible; SCM, set covering machine; RF-C, random forest classification; RF-mC, random forest multi-class classification; RF-R, random forest regression

^1^Indicates the dataset from which training and testing sets were derived.
