## Supplemental Table 4 for "Evaluation of parameters affecting performance and reliability of machine learning-based antibiotic susceptibility testing from whole genome sequencing data"

**Table S4.** Performance (mean with 95% confidence intervals) of predictive models for ciprofloxacin resistance from the primary set covering machine and random forest analyses.

|  |  | N. gonorrhoeae (aggregate dataset) | N. gonorrhoeae (1) | N. gonorrhoeae (2) | N. gonorrhoeae (3) | N. gonorrhoeae (4) | N. gonorrhoeae (5) | N. gonorrhoeae (6) | N. gonorrhoeae (7) | K. pneumoniae (8) | A. baumannii (9) |
| --- | --- | --- | --- | --- | --- | --- | --- | --- | --- | --- | --- |
|  | N_genomes_ | 3946 | 886 | 1102 | 672 | 383 | 714 | 204 | 398 | 1560 | 702 |
|  | N_S_ (EUCAST) | 2131 | 659 | 500 | 292 | 181 | 372 | 30 | 271 | N/A | 96 |
|  | N_NS_ (EUCAST) | 1815 | 227 | 602 | 379 | 202 | 342 | 174 | 127 | N/A | 606 |
|  | N_S_ (CLSI) | 2154 | 664 | 503 | 303 | 190 | 374 | 30 | 272 | 197 | 96 |
|  | N_NS_ (CLSI) | 1792 | 222 | 599 | 368 | 193 | 340 | 174 | 126 | 1363 | 606 |
|  | N_kmers_ | 7.21E+06 | 3.43E+06 | 3.41E+06 | 4.89E+06 | 4.58E+06 | 4.41E+06 | 2.75E+06 | 3.15E+06 | 5.42E+07 | 2.52E+07 |
| SCM (S/NS) | 5-CV bACC^1^  (EUCAST) | 98.76  (98.64-98.88) | 97.86  (97.37-98.34) | 99.18  (99.03-99.33) | 98.28  (98.02-98.54) | 97.53  (96.77-98.29) | 98.57  (98.39-98.74) | 99.53  (99.36-99.7) | 98.96  (98.68-99.25) | N/A | 97.29  (96.92-97.66) |
|  | bACC (EUCAST) | 98.41  (98.25-98.58) | 98.03  (97.12-98.94) | 99  (98.7-99.29) | 97.80  (97.32-98.29) | 95.99  (94.19-97.8) | 99.39  (99.14-99.63) | 98.75  (97.48-100) | 97.85  (97.08-98.61) | N/A | 97.83  (96.67-98.99) |
|  | Sensitivity (EUCAST) | 97.73 (97.41-98.04) | 97 (95.41-98.59) | 98.71  (98.26-99.16) | 97.12 (96.09-98.16) | 94.79 (92.78-96.8) | 99.49 (99.08-99.91) | 98.33 (97.27-99.39) | 96.79 (95.21-98.37) | N/A | 98.42 (97.88-98.96) |
|  | Specificity (EUCAST) | 99.10 (98.89-99.31) | 99.06 (99.41-98.71) | 99.28  (99.01-99.55) | 98.48 (97.71-99.25) | 97.19 (94.57-99.82) | 99.28 (98.72-99.84) | 99.17 (97.28-101.1) | 98.91 (98.27-99.55) | N/A | 97.24 (95.11-99.36) |
|  | 5-CV bACC^1^  (CLSI) | 99.09  (99.02-99.15) | 98.13  (97.81-98.46) | 99.02  (98.88-99.15) | 99.31  (99.12-99.49) | 100  (100-100) | 99.68  (99.58-99.78) | 99.53  (99.36-99.7) | 98.65  (98.4-98.9) | 97.22  (96.78-97.65) | 97.29  (96.92-97.66) |
|  | bACC (CLSI) | 98.80 (98.67-98.93) | 98.13 (97.45-98.8) | 98.95  (98.68-99.22) | 99.32 (99.70-98.95) | 99.54 (99-100.1) | 99.52 (99.41-99.62) | 98.75 (97.48-100) | 99.17 (98.67-99.68) | 90.73 (88.37-93.08) | 97.83  (96.67-98.99) |
|  | Sensitivity (CLSI) | 98.82 (98.62-99.01) | 97.64 (96.37-98.9) | 98.86  (98.43-99.29) | 99.36 (98.91-99.81) | 99.55 (98.54-100.6) | 99.58 (99.25-99.9) | 98.33 (97.27-99.39) | 99.02 (98.12-99.93) | 95.01 (94.30-95.73) | 98.42 (97.88-98.96) |
|  | Specificity (CLSI) | 98.78 (98.61-98.96) | 98.62 (98.33-98.91) | 99.04  (98.52-99.55) | 99.29 (98.8-99.77) | 99.54 (99-100.1) | 99.46 (99.01-99.91) | 99.17 (97.28-101.1) | 99.32 (98.75-99.9) | 86.44 (81.25-91.63) | 97.24 (95.11-99.36) |
| RF-C (S/NS) | OOB bACC^1^  (EUCAST) | 98.66  (98.28-99.04) | 98.54  (97.98-99.11) | 99.4  (99.11-99.69) | 97.03  (96.49-97.57) | 96.05  (95.68-96.42) | 98.76  (98.42-99.09) | 99.51  (99.07-99.94) | 97.92  (97.4-98.44) | N/A | 91.73  (90.66-92.81) |
|  | bACC (EUCAST) | 98.41 (98.15-98.67) | 98.35  (98.24-98.47) | 99.32  (99.22-99.42) | 96.96  (96.86-97.06) | 95.87  (95.77-95.98) | 98.53  (98.43-98.64) | 99.62  (99.55-99.69) | 97.88  (97.77-97.98) | N/A | 91.62  (90.39-92.85) |
|  | Sensitivity (EUCAST) | 98.54 (97.93-99.15) | 97.09  (96.7-97.48) | 99.4  (99.08-99.72) | 96.54  (96.28-96.8) | 94.21  (94-94.42) | 98.33  (98.15-98.5) | 99.3  (99.13-99.47) | 97.29  (97.09-97.49) | N/A | 97.69  (97.02-98.36) |
|  | Specificity (EUCAST) | 98.29 (97.88-98.69) | 99.62  (99.32-99.92) | 99.24  (98.9-99.58) | 97.39  (97.11-97.66) | 97.54  (97.32-97.76) | 98.74  (98.56-98.92) | 99.94  (99.88-100) | 98.47  (98.28-98.65) | N/A | 85.56 (83.33-87.78) |
|  | OOB bACC^1^  (CLSI) | 98.88  (98.42-99.35) | 98.44  (98.21-98.66) | 99.7  (99.32-100.1) | 98.24  (97.9-98.58) | 99.07  (98.69-99.45) | 99.45  (98.99-99.92) | 99.51  (99.07-99.94) | 98.57  (98.23-98.9) | 89.90  (87.78-92.02) | 91.73  (90.66-92.81) |
|  | bACC (CLSI) | 98.76 (98.29-99.24) | 98.48  (98.31-98.64) | 99.5  (99.37-99.62) | 98.08  (97.92-98.24) | 98.83  (98.7-98.96) | 99.34  (99.18-99.5) | 99.78  (99.7-99.87) | 99.62  (99.55-99.69) | 91.42 (90.16-92.68) | 91.62  (90.39-92.85) |
|  | Sensitivity (CLSI) | 98.76 (98.29-99.24) | 99.15  (98.76-99.55) | 99.39  (99.06-99.73) | 97.17  (96.87-97.47) | 97.94  (97.67-98.2) | 99.38  (99.13-99.63) | 99.3  (99.13-99.47) | 98.01  (97.72-98.31) | 86.14 (83.91-88.37) | 97.69  (97.02-98.36) |
|  | Specificity (CLSI) | 98.95 (98.21-99.68) | 97.8  (97.4-98.19) | 99.6  (99.35-99.85) | 98.99  (98.71-99.27) | 99.72  (99.54-99.91) | 99.3  (99.09-99.51) | 99.94  (99.88-100) | 99.1  (98.86-99.34) | 96.70 (95.40-98.01) | 85.56 (83.33-87.78) |
| RF-mC (MIC) | OOB 1-tier accuracy^1^ | 91.56  (91.25-91.88) | 88.27 (85.09-91.45) | 94.53 (92.02-97.04) | 93.43 (90.49-96.36) | 95.77 (93.36-98.18) | 82.68 (80.37-85) | 83.56 (81.64-85.47) | 95.52 (92.82-98.22) |  |  |
|  | Exact match | 73.99  (73.29-74.7) | 80.8 (78.6-83) | 67.83 (66.35-69.31) | 71.96 (70.33-73.58) | 77.58 (75.5-79.66) | 63.85 (61.19-66.5) | 72.31 (69.59-75.04) | 86.26 (84.44-88.08) | N/A | N/A |
|  | 1-tier accuracy | 91.51  (91.23-91.79) | 87.36 (84.92-89.8) | 95.52 (93.93-97.1) | 93.26 (91.24-95.29) | 94.33 (91.2-97.46) | 82.4 (80.83-83.98) | 83.94 (82.78-85.1) | 95.1 (92.36-97.84) | N/A | N/A |
|  | bACC^2^ (EUCAST) | 96.79  (96.56-97.02) | 96.76 (94.52-99) | 98.62 (98.06-99.18) | 95.98 (94.8-97.15) | 95.03 (92.99-97.07) | 97.74 (96.14-99.34) | 99.42 (98.95-99.89) | 94.8 (92.66-96.94) | N/A | N/A |
|  | Sensitivity (EUCAST) | 99.1  (98.87-99.32) | 94.84 (91.85-97.83) | 97.57 (96.72-98.43) | 94.28 (92.78-95.78) | 92.46 (90.84-94.09) | 96.9 (95.23-98.58) | 98.83 (98.08-99.58) | 91.17 (89.01-93.34) | N/A | N/A |
|  | Specificity (EUCAST) | 97.94  (97.81-98.08) | 98.68 (97.15-100.2) | 99.66 (99.35-99.98) | 97.67 (96.67-98.68) | 97.59 (95.05-100.1) | 98.58 (96.95-100.2) | 100 (99.7-100.3) | 98.42 (96.26-100.6) | N/A | N/A |
|  | bACC^2^ (CLSI) | 97.07  (96.69-97.44) | 96.57 (94.41-98.74) | 98.28 (96.67-99.9) | 97.61 (96.23-99) | 98.26 (96.63-99.9) | 97.8 (96.39-99.21) | 99.42 (98.95-99.89) | 94.83 (92.95-96.7) | N/A | N/A |
|  | Sensitivity (CLSI) | 99.24  (99.16-99.32) | 94.84 (91.95-97.74) | 97.46 (95.77-99.14) | 95.98 (93.74-98.22) | 97.95 (96.54-99.37) | 96.29 (93.78-98.8) | 98.83 (98.08-99.58) | 91.66 (89.24-94.07) | N/A | N/A |
|  | Specificity (CLSI) | 98.15  (97.97-98.33) | 98.3 (96.82-99.79) | 99.11 (97.45-100.8) | 99.25 (98.61-99.9) | 98.57 (96.53-100.6) | 99.31 (98.85-99.77) | 100 (99.7-100.3) | 98 (96.42-99.57) | N/A | N/A |
| RF-R (log_2_MIC) | OOB 1-tier accuracy^1^ | 87.68 (86.87-88.5) | 91.56 (90.72-92.4) | 87.56 (86.67-88.44) | 88.38 (87.82-88.93) | 90.38 (89.41-91.35) | 79.82 (78.92-80.72) | 84.95 (84.16-85.74) | 89.41 (88.87-89.96) |  |  |
|  | Exact match | 67.63 (66.89-68.36) | 73.65 (73-74.31) | 69.29 (68.72-69.86) | 67.27 (66.91-67.64) | 71.66 (70.94-72.39) | 54.08 (53.39-54.78) | 64.81 (63.78-65.83) | 78.48 (77.62-79.34) | N/A | N/A |
|  | 1-tier accuracy | 84.38 (83.57-85.19) | 87.32 (86.51-88.13) | 84.8 (83.71-85.89) | 85.59 (84.88-86.3) | 87.49 (86.64-88.33) | 76.36 (75.93-76.78) | 80.92 (79.99-81.84) | 85.82 (85.07-86.58) | N/A | N/A |
|  | bACC^2^ (EUCAST) | 95.96 (95.2-96.73) | 93.92 (93.14-94.71) | 96.39 (95.63-97.14) | 95.19 (94.36-96.02) | 97.39 (96.72-98.06) | 94.86 (94.03-95.69) | 93.52  (92.11-94.93) | 93.09  (91.85-94.33) | N/A | N/A |
|  | Sensitivity (EUCAST) | 96.26 (95.47-97.05) | 91.19 (90.45-91.93) | 96 (95.25-96.75) | 94.53 (93.51-95.56) | 95.3 (94.24-96.37) | 95.85 (95.02-96.68) | 95.7  (94.03-97.36) | 87.94  (86.48-89.4) | N/A | N/A |
|  | Specificity (EUCAST) | 95.67 (94.9-96.44) | 96.66 (95.76-97.56) | 96.78 (96.01-97.55) | 95.85 (95.19-96.51) | 99.47 (99.07-99.88) | 93.87 (92.98-94.76) | 91.34  (89.55-93.13) | 98.24  (97.16-99.32) | N/A | N/A |
|  | bACC^2^ (CLSI) | 97.68 (96.92-98.45) | 96.72 (95.89-97.55) | 97.48 (96.69-98.27) | 95.85 (95.31-96.39) | 96.54 (95.9-97.18) | 97.79 (97.07-98.51) | 94.28  (93.23-95.33) | 97.69  (96.52-98.86) | N/A | N/A |
|  | Sensitivity (CLSI) | 98.09 (97.42-98.75) | 96.47 (95.73-97.21) | 98.07 (97.44-98.7) | 98.06 (97.62-98.5) | 98.04 (97.56-98.52) | 98.67 (98.02-99.32) | 96.4  (95.34-97.47) | 97.81  (96.26-99.36) | N/A | N/A |
|  | Specificity (CLSI) | 97.28 (96.31-98.25) | 96.97 (95.97-97.97) | 96.9 (95.91-97.89) | 93.64 (92.86-94.43) | 95.04 (94.19-95.89) | 96.92 (96.09-97.75) | 92.16  (91.07-93.26) | 97.58  (96.58-98.58) | N/A | N/A |

S, susceptible; NS, non-susceptible; SCM, set covering machine; RF-C, random forest classification; RF-mC, random forest multi-class classification; RF-R, random forest regression; 5-CV, five-fold cross-validation; OOB, out-of-bag estimate

^1^bACC (for SCM and RF-C) or 1-tier accuracy (for RF-mC and RF-R) based on five-fold cross-validation (for SCM) or bootstrap aggregating (for RF-C, RF-mC, and RF-R) of training sets.

^2^bACC based on categorical agreement (by either the EUCAST or CLSI breakpoint) between phenotypic and predicted MICs.
