## Supplemental Table 5 for "Evaluation of parameters affecting performance and reliability of machine learning-based antibiotic susceptibility testing from whole genome sequencing data"

**Table S5.** Performance (mean with 95% confidence intervals) of predictive models for azithromycin resistance from the primary set covering machine and random forest analyses.

|  |  | N. gonorrhoeae (aggregate dataset) | N. gonorrhoeae (1) | N. gonorrhoeae (2) | N. gonorrhoeae (3) | N. gonorrhoeae (4) | N. gonorrhoeae (5) | N. gonorrhoeae (6) | N. gonorrhoeae (7) |
| --- | --- | --- | --- | --- | --- | --- | --- | --- | --- |
|  | N_genomes_ | 3946 | 886 | 1102 | 671 | 383 | 714 | 204 | 398 |
|  | N_S_ (EUCAST) | 2119 | 656 | 192 | 232 | 103 | 496 | 131 | 355 |
|  | N_NS_ (EUCAST) | 1827 | 230 | 910 | 439 | 280 | 218 | 73 | 43 |
|  | N_S_ (CLSI) | 3410 | 876^1^ | 807 | 440 | 175 | 701^1^ | 201^1^ | 396^1^ |
|  | N_NS_ (CLSI) | 536 | 10^1^ | 295 | 231 | 208 | 13^1^ | 3^1^ | 2^1^ |
|  | N_kmers_ | 7.21E+06 | 3.43E+06 | 3.41E+06 | 4.89E+06 | 4.58E+06 | 4.41E+06 | 2.75E+06 | 3.15E+06 |
| SCM (S/NS) | 5-CV bACC^1^  (EUCAST) | 79.11  (78.9-79.31) | 74.15  (71.37-76.92) | 67.02  (62.02-72.02) | 83.99  (81.75-86.22) | 78.87  (75.53-82.21) | 65.81  (60.97-70.65) | 83.81  (79.83-87.79) | 78.03  (72.49-83.57) |
|  | bACC (EUCAST) | 77.6 (76.74-78.46) | 69.69 (66.79-72.59) | 62.44 (57.86-67.02) | 73.59 (70.95-76.23) | 72.68 (69.84-75.52) | 64.73 (62.96-66.5) | 68.78 (63.52-74.04) | 69.6 (66.22-72.97) |
|  | Sensitivity (EUCAST) | 85.39 (81.96-88.82) | 49.76 (41.05-58.47) | 96.94 (95.55-98.33) | 88.93 (85.43-92.42) | 97.18 (95.33-99.02) | 42.08 (32.26-51.9) | 53.35 (42.2-64.49) | 41.12 (34.19-48.05) |
|  | Specificity (EUCAST) | 69.82 (67.79-71.85) | 89.62 (85.43-93.81) | 27.94  (17.51-38.37) | 58.26 (50.12-66.39) | 48.18 (42.11-54.24) | 87.38 (79.92-94.84) | 84.21 (76.53-91.89) | 98.07 (96.97-99.18) |
|  | 5-CV bACC^1^  (CLSI) | 87.75  (86.35-89.15) | N/A | 87.72  (85.79-89.65) | 94.63  (93.83-95.44) | 94.87  (93.42-96.33) | N/A | N/A | N/A |
|  | bACC (CLSI) | 85.32 (83.68-86.95) | N/A | 81.46 (79.79-83.13) | 92.13 (89.99-94.27) | 88.42 (87.06-89.78) | N/A | N/A | N/A |
|  | Sensitivity (CLSI) | 71.6  (68.07-75.13) | N/A | 66.04  (62.12-69.96) | 90.29 (84.86-95.72) | 86.64 (83.23-90.04) | N/A | N/A | N/A |
|  | Specificity (CLSI) | 99.03  (98.7-99.37) | N/A | 96.88 (95.01-98.75) | 93.97 (91.63-96.31) | 90.21 (85.88-94.53) | N/A | N/A | N/A |
| RF-C (S/NS) | OOB bACC^1^  (EUCAST) | 84.57  (83.57-85.56) | 78.32  (76.26-80.37) | 67.34  (66.08-68.60) | 82.34  (80.50-84.17) | 82.67  (81.81-83.53) | 75.54  (73.53-77.54) | 80.42  (78.97-81.88) | 78.55  (77.21-79.90) |
|  | bACC (EUCAST) | 84.04  (83.11-84.98) | 73.01  (71.85-74.18) | 65.16  (64.10-66.22) | 81.05  (79.38-82.72) | 79.07  (76.94-81.19) | 71.96 (70.80-73.13) | 78.65  (76.86-80.45) | 75.5  (74.15-76.85) |
|  | Sensitivity (EUCAST) | 84.36  (82.68-86.03) | 57.4  (56.36-58.43) | 95.15 (93.47-96.82) | 90.06  (87.67-92.45) | 93.08  (90.74-95.41) | 56.53 (53.39-59.68) | 69.23  (65.72-72.74) | 51.73  (49.24-54.22) |
|  | Specificity (EUCAST) | 83.73  82.16-85.31) | 88.63  (87.15-90.11) | 35.16 (31.82-38.51) | 72.04  (70.84-73.23) | 65.06  (62.41-67.71) | 87.39 (84.84-89.94) | 88.08  (86.06-90.10) | 99.27  (98.28-100.3) |
|  | OOB bACC^1^  (CLSI) | 89.66  (88.64-90.67) | N/A | 82.81  (81.78-83.84) | 94.09  (93.15-95.03) | 87.61  (86.22-89) | N/A | N/A | N/A |
|  | bACC (CLSI) | 88.03  (86.28-89.79) | N/A | 82.34 (80.89-83.80) | 93.42  (92.41-94.44) | 85.37  (83.69-87.06) | N/A | N/A | N/A |
|  | Sensitivity (CLSI) | 77.05  (73.46-80.63) | N/A | 72.89 (68.78-77.01) | 89.89  (88.86-90.91) | 85.09  (82.92-87.26) | N/A | N/A | N/A |
|  | Specificity (CLSI) | 99.02  (98.74-99.30) | N/A | 91.79 (89.77-93.81) | 96.96  (95.84-98.09) | 85.66  (82.19-89.13) | N/A | N/A | N/A |
| RF-mC (MIC) | OOB 1-tier accuracy^1^ | 84.06  (82.9-85.23) | 75.82  (74.21-77.43) | 87.57  (85.15-89.98) | 87.28  (85.82-88.74) | 79.8  (78.4-81.19) | 88  (86.2-89.81) | 92.9  (91.43-94.38) | 95.3  (94.13-96.47) |
|  | Exact match | 48.18  (47.47-48.89) | 24.39 (22.51-26.27) | 50.81 (49.4-52.21) | 59.54 (57.23-61.84) | 58.1 (56.72-59.49) | 36.01 (33.75-38.26) | 41.12 (39.23-43.01) | 59.15 (57.88-60.41) |
|  | 1-tier accuracy | 83.3  (82.94-83.67) | 73.4 (71.52-75.28) | 85.55 (83.61-87.5) | 85.18 (83.42-86.94) | 76.89 (75.4-78.37) | 85.01 (83.19-86.83) | 90.71 (88.47-92.94) | 93.66 (91.47-95.84) |
|  | bACC^3^ (EUCAST) | 82.36  (81.75-82.98) | 72.93 (72.16-73.71) | 64.94 (64.11-65.78) | 81.43 (80.83-82.04) | 80.76 (79.36-82.17) | 73.44 (71.98-74.9) | 76.16 (74.84-77.49) | 76.41 (75.4-77.43) |
|  | Sensitivity (EUCAST) | 84.11  (83.83-84.39) | 61.12 (60.15-62.09) | 96.16 (94.99-97.33) | 89.04 (87.98-90.09) | 89.36 (87.89-90.83) | 60.53 (58-63.06) | 65.94 (64.77-67.11) | 53.61 (52.15-55.07) |
|  | Specificity (EUCAST) | 80.62  (79.52-81.71) | 84.75 (83.64-85.85) | 33.73 (32.73-34.72) | 73.83 (73.07-74.58) | 72.17 (69.92-74.41) | 86.35 (85.5-87.2) | 86.39 (84.29-88.49) | 99.22 (98.58-99.86) |
|  | bACC^3^ (CLSI) | 87.79  (85.92-89.67) | N/A | 82.85 (80.03-85.66) | 90.08 (88.23-91.93) | 85.97 (84.75-87.2) | N/A | N/A | N/A |
|  | Sensitivity (CLSI) | 80.71  (79.14-82.29) | N/A | 74.67 (72.92-76.41) | 86.19 (84.7-87.67) | 87.73 (85.49-89.96) | N/A | N/A | N/A |
|  | Specificity (CLSI) | 95.1  (92.89-97.3) | N/A | 93.13 (92.35-93.92) | 95.15 (94.33-95.97) | 84.23 (83.54-84.92) | N/A | N/A | N/A |
| RF-R (log_2_MIC) | OOB 1-tier accuracy^1^ | 79.66  (77.97-81.35) | 77.88  (75.63-80.13) | 79.35  (78.19-80.51) | 78.57  (76.24-80.89) | 78.85  (77.52-80.17) | 83.21  (81.03-85.38) | 87.52  (85.52-89.52) | 85.61  (84.47-86.75) |
|  | Exact match | 52.89 (49.27-56.52) | 44.63 (42.38-46.87) | 52.37 (49.74-55) | 54.31 (51.28-57.34) | 52.3 (51.25-53.36) | 53.66 (52.09-55.22) | 64.08 (62.2-65.96) | 64.58 (62.45-66.7) |
|  | 1-tier accuracy | 77.23 (73.77-80.7) | 72.75 (70.67-74.82) | 76.06 (73.3-78.82) | 75.91 (73.07-78.74) | 74.91 (72.78-77.04) | 78.11 (76.6-79.63) | 81.81 (79.17-84.44) | 83.34 (81.16-85.53) |
|  | bACC^3^ (EUCAST) | 79.43  (78.48-80.39) | 76.38 (74.3-78.46) | 57.1 (54.49-59.72) | 69.6 (67.39-71.81) | 71.12 (69.26-72.99) | 77.37 (75.52-79.21) | 75.86 (73.97-77.75) | 84.66 (82.57-86.75) |
|  | Sensitivity (EUCAST) | 90.88  (89.41-92.34) | 67.3 (65.03-69.56) | 99.13 (96.62-101.6) | 97.9 (94.97-100.8) | 97.84 (96.12-99.55) | 79.53 (78.31-80.74) | 77.34 (75.68-79) | 82.73 (81.18-84.28) |
|  | Specificity (EUCAST) | 67.99  (67.28-68.7) | 85.47 (83.54-87.4) | 15.08 (12.31-17.84) | 41.3 (39.61-42.99) | 44.41 (42.37-46.46) | 75.2 (72.56-77.85) | 74.38 (72.08-76.68) | 86.59 (83.63-89.55) |
|  | bACC^3^ (CLSI) | 86.58 (84.94-88.22) | N/A | 84.44 (81.71-87.17) | 94.18 (92.55-95.82) | 87.23 (85.77-88.7) | N/A | N/A | N/A |
|  | Sensitivity (CLSI) | 73.78 (71.58-75.98) | N/A | 79.8 (77.09-82.51) | 93.98 (92.97-94.99) | 90.02 (88.04-92.01) | N/A | N/A | N/A |
|  | Specificity (CLSI) | 99.38 (97.98-100.8) | N/A | 89.08 (86.26-91.89) | 94.39 (92.08-96.7) | 84.44 (83.01-85.87) | N/A | N/A | N/A |

S, susceptible; NS, non-susceptible; SCM, set covering machine; RF-C, random forest classification; RF-mC, random forest multi-class classification; RF-R, random forest regression; 5-CV, five-fold cross-validation; OOB, out-of-bag estimate

^1^Azithromycin NS by the CLSI breakpoint was not assessed for dataset 1 or 5-7 due to low representation (<15) of NS strains.

^1^bACC (for SCM and RF-C) or 1-tier accuracy (for RF-mC and RF-R) based on five-fold cross-validation (for SCM) or bootstrap aggregating (for RF-C, RF-mC, and RF-R) of training sets.

^3^bACC based on categorical agreement (by either the EUCAST or CLSI breakpoint) between phenotypic and predicted MICs.
