## Supplemental Table 6 for "Evaluation of parameters affecting performance and reliability of machine learning-based antibiotic susceptibility testing from whole genome sequencing data"

**Table S6.** Summary of approach for the additional classification analyses.

| Testing set | Training set | AZM resistance metric | Classification method | Results |
| --- | --- | --- | --- | --- |
| Subsamples from aggregate GC dataset with strains with AZM MICs from 0.06-1 μg/mL removed | Subsamples from aggregate GC dataset with strains with AZM MICs from 0.06-1 μg/mL removed | AZM NS (EUCAST,  >0.25 μg/mL) | SCM/RF-C | Fig S1c |
| Subsamples from aggregate GC dataset with strains with AZM MICs from 0.25-4 μg/mL removed | Subsamples from aggregate GC dataset with strains with AZM MICs from 0.06-1 μg/mL removed | AZM NS (CLSI,  >1 μg/mL) | SCM/RF-C | Fig S1c |
| Subsamples from datasets 1-7 | Subsamples from aggregate GC dataset | AZM NS (EUCAST,  >0.25 μg/mL) | RF-C | Fig 2a-b |
|  | Subsamples from aggregate GC dataset excluding isolates from dataset of testing isolates |  |  |  |
| Subsamples from dataset 2 after down-sampling to match sample size and MIC distribution of dataset 4 | Subsamples from dataset 2 after down-sampling to match sample size and MIC distribution of dataset 4 | AZM NS (EUCAST,  >0.25 μg/mL) | SCM/RF-C | Fig S2c |
| Subsamples from dataset 4 after down-sampling to match sample size MIC distribution of dataset 2 | Subsamples from dataset 4 after down-sampling to match sample size MIC distribution of dataset 2 | AZM NS (EUCAST,  >0.25 μg/mL) | RF-C | Fig S2c |
| Subsamples from the *K. pneumoniae* dataset after down-sampling to equalize the number of S and NS strains | Subsamples from the *K. pneumoniae* dataset after down-sampling to equalize the number of S and NS strains | CIP NS (CLSI, >1 μg/mL) | SCM/RF-C | Fig S3c-d |
| Subsamples from the *A. baumannii* dataset after down-sampling to equalize the number of S and NS strains | Subsamples from the *A. baumannii* dataset after down-sampling to equalize the number of S and NS strains | CIP NS (EUCAST/CLSI, >1 μg/mL) | SCM/RF-C | Fig S3c-d |

GC, gonococcal; AZM, azithromycin; MICs, minimum inhibitory concentrations; NS, non-susceptible; SCM, set covering machine; RF-C, random forest classification
