## Supplementary figures and images for "Evaluation of parameters affecting performance and reliability of machine learning-based antibiotic susceptibility testing from whole genome sequencing data"

### Supplemental Figure 1

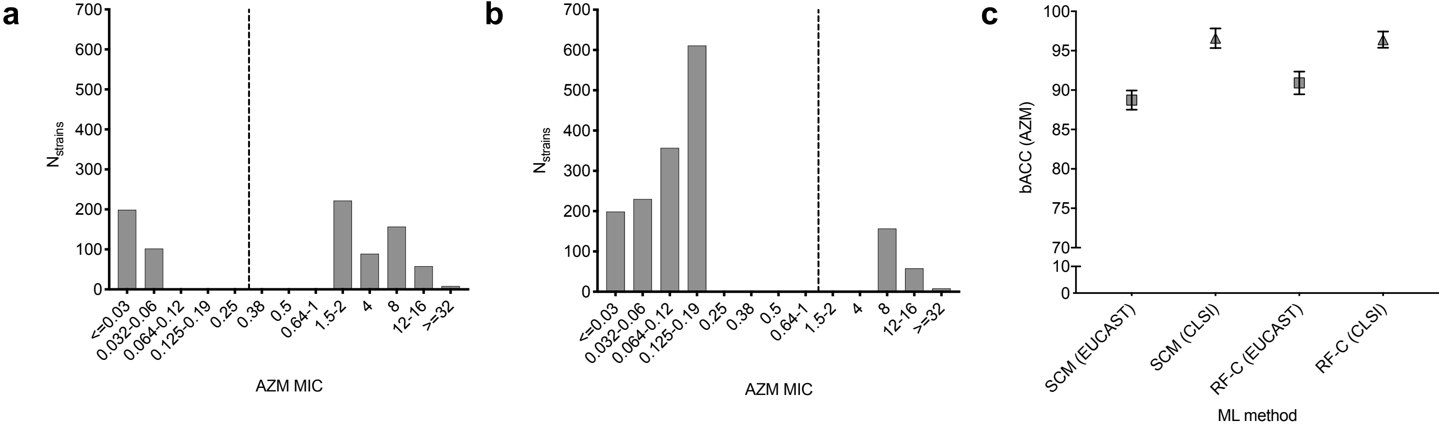

### Supplemental Figure 2

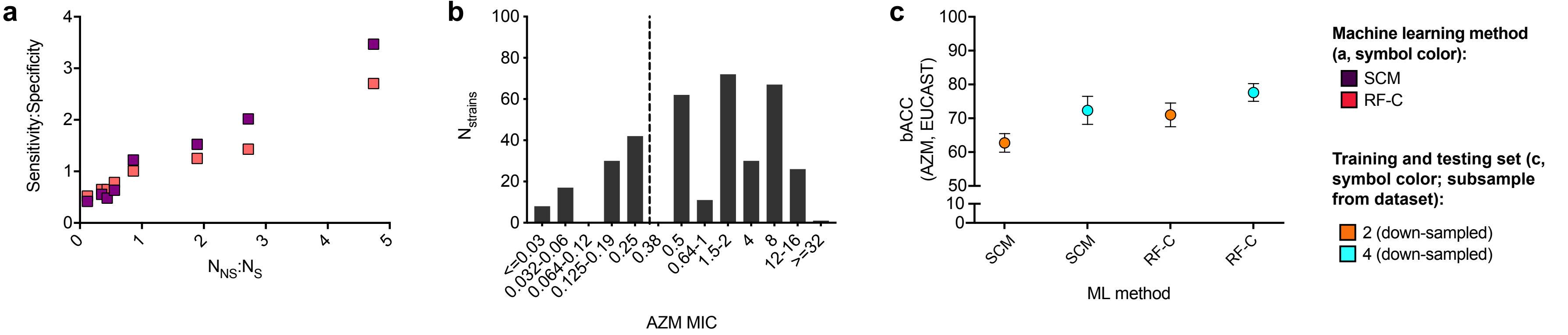

### Supplemental Figure 3

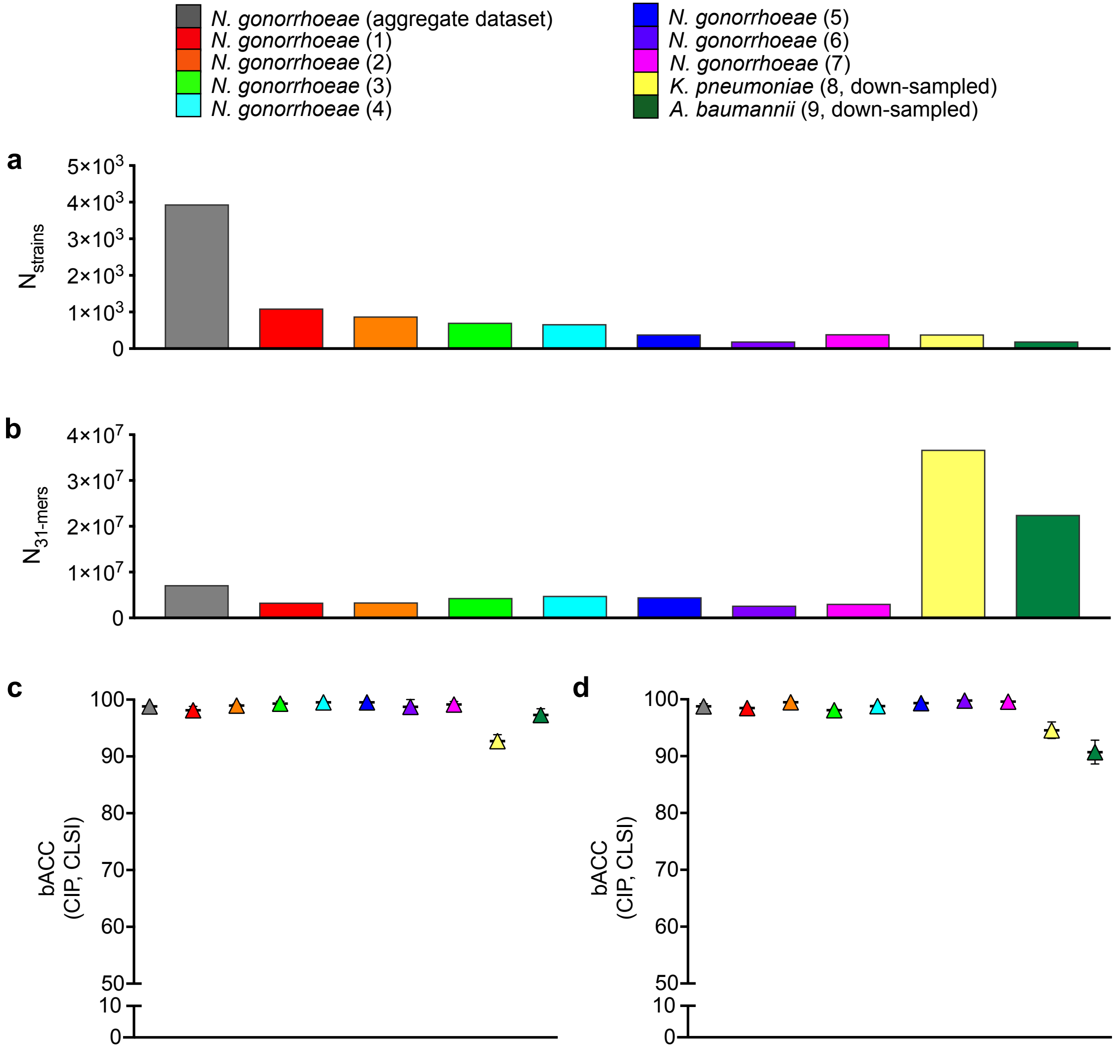
